# Widespread epistasis shapes RNA Polymerase II active site function and evolution

**DOI:** 10.1101/2023.02.27.530048

**Authors:** Bingbing Duan, Chenxi Qiu, Sing-Hoi Sze, Craig Kaplan

## Abstract

Multi-subunit RNA Polymerases (msRNAPs) are responsible for transcription in all kingdoms of life. These enzymes rely on dynamic, highly conserved active site domains such as the so-called “trigger loop” (TL) to accomplish steps in the transcription cycle. Mutations in the RNA polymerase II (Pol II) TL confer a spectrum of biochemical and genetic phenotypes that suggest two main classes, which decrease or increase catalysis or other nucleotide addition cycle (NAC) events. The Pol II active site relies on networks of residue interactions to function, and mutations likely perturb these networks in ways that may alter mechanisms. We have undertaken a structural genetics approach to reveal residue interactions within and surrounding the Pol II TL – determining its “interaction landscape” – by deep mutational scanning in *Saccharomyces cerevisiae* Pol II. This analysis reveals connections between TL residues and surrounding domains, demonstrating that TL function is tightly coupled to its specific enzyme context.

## INTRODUCTION

Transcription from cellular genomes is performed by conserved multi-subunit RNA polymerases (msRNAPs)^1–3^. While bacteria and archaea use a single msRNAP to transcribe all RNAs^4–6^, eukaryotes have at least three msRNAPs (Pol I, II, and III) for different RNA types^7–10^. RNA synthesis by msRNAPs occurs by iterative nucleotide addition cycles (NAC) of nucleotide selection, catalysis and polymerase translocation^11–15^. The msRNAPs active sites accomplish these NAC steps using two conformationally flexible domains, the bridge helix (BH) and trigger loop (TL)^5,12,16–19^. The multi-functional natures of the BH and TL likely underlie their striking conservation, serving as interesting models for studying the function and evolution of extremely constrained protein domains.

Most catalytic cycle events involve concerted conformational changes in the TL, and potentially the BH^12–14,19–23^. The extent to which these conformations are supported by, or communicate with, the surrounding enzymatic context can be examined by structural or computational approaches, but functional importance needs to be determined experimentally. The BH is straight in most msRNAP structures^7,8,12,13,24^, but appears kinked in *Thermus thermophilus* (bacteria) RNAP^5,25^. Simulations suggest these dynamics may promote msRNAP translocation^16,25–28^. The TL adopts various conformations to support its multiple functions. During each NAC, the TL nucleotide interaction region (NIR) discriminates correct NTPs from mismatched ones, initiating a shift from a catalytic-disfavoring “open” state to a catalytic-favoring “closed” state^11,20,29,30^. This closure promotes catalysis^20,31^, and subsequent TL reopening is proposed to support polymerase translocation to the next template DNA position, allowing for the next NAC^32–34^(**Fig. 1a**). Disrupting the TL with mutations affects transcription at all stages^13,17,23,35–39^. For instance, NIR mutations impair TL-substrate interactions, resulting in reduced catalysis and elongation rate in vitro (Loss of function, LOF)^17,23,36,37,40,41^. Mutations in the TL hinge and C-terminal regions appear to shift the TL toward the active state, enhancing catalysis and elongation rate but impairing transcription fidelity (Gain of function, GOF)^17,42,43^. Mutations may also affect other transcription processes such as proofreading, pausing and termination^12,43–46^, as TL confirmations are associated with these activities in addition to catalysis and translocation.

**Figure 1.**
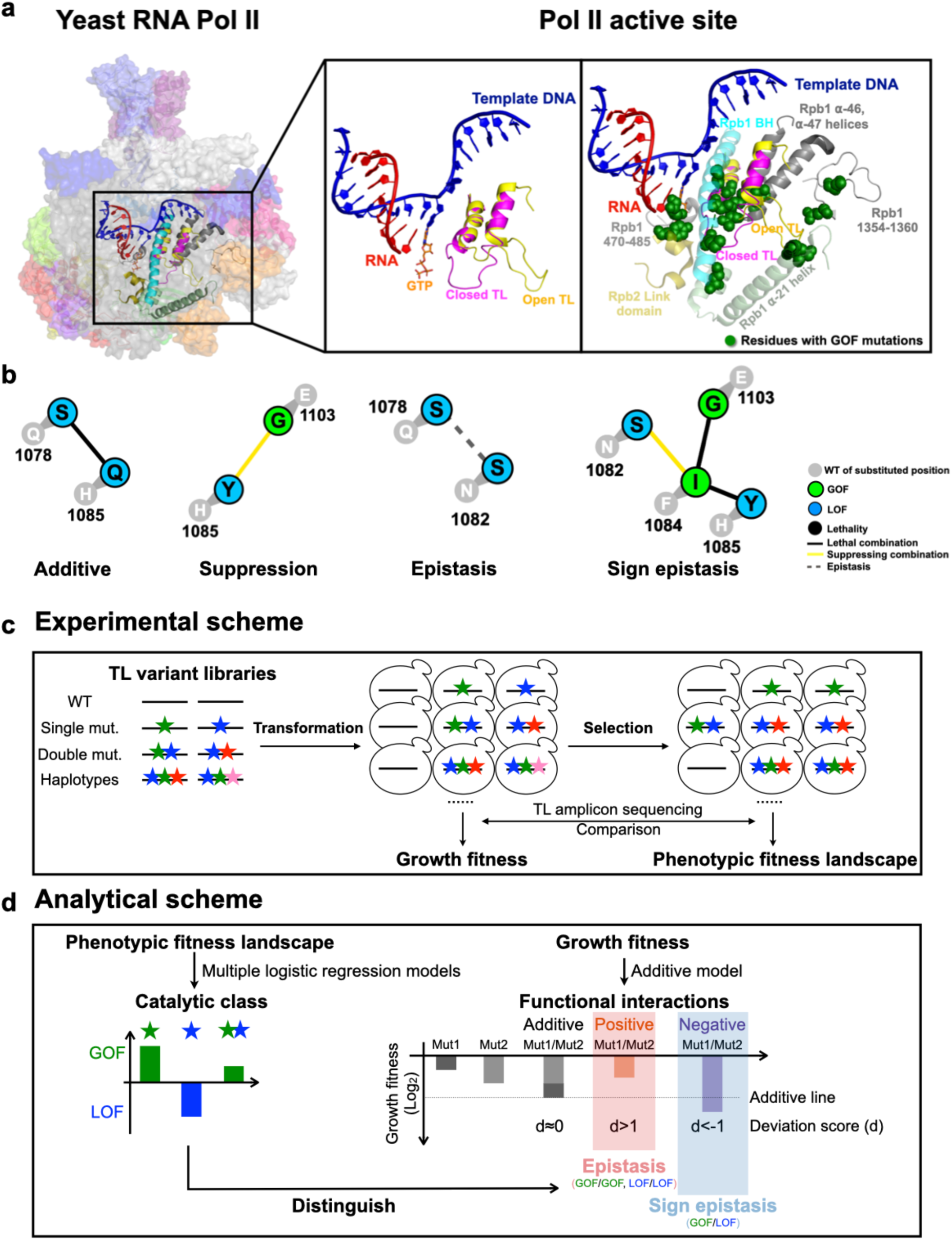
Schematics of the Pol II active site interaction landscape. **a.** The Pol II active site is embedded in the center of a 12-subunit complex (left panel). Pol II functions are supported by distinct TL conformational states. An open TL (PDB: 5C4X)^42^ and closed TL (PDB: 2E2H)^12^ conformations are shown in the middle panel. GOF mutations have been identified in the TL and its proximal domains (right panel), suggesting TL mobility and function may be impacted by adjacent residues. **b.** Examples of inter-residue genetic interactions. WT residues are shown in grey circles with number indicating residue position in Rpb1. Mutant substitutions are shown in colored circles, with color representing mutant class. Colored lines between mutant substitutions represent types of genetic interactions. **c.** Overview of experimental approach. We synthesized 10 libraries of TL variants represented by colored stars. Libraries were transformed into WT or mutated yeast strains. A selection assay was subsequently performed by scraping and replating the transformants onto different media for phenotyping. DNA was extracted from yeast from all conditions, and went through TL region amplification, and Illumina sequencing. Read counts for variants on general conditions were used to determine growth fitness, while read counts on other conditions were used to determine the phenotypic fitness landscape (see **Methods**). **d.** Overview of analytical approach for determining interaction landscape. Mutant conditional growth fitnesses were calculated using allele frequencies under selective growth conditions and subjected to two logistic regression models for classification/prediction of catalytic defects. Double mutant interactions were computed using growth fitness. Classification allowed epistatic interactions to be deduced from double mutant growth fitness (see **Methods**).

All TL conformational dynamics and functions are likely balanced by residue interactions within and around the TL^41,45,47–49^. Embedded in the conserved active site, the TL interacts with surrounding domains such as the BH and ⍺46-47 helices, forming a five-helix bundle enclosing a hydrophobic pocket that supports these dynamics **(Fig. 1a)**^12,42^. Mutations in TL adjacent domains (e.g., BH GOF T834P and LOF T834A, funnel helix ⍺-21 GOF S713P, and Rpb2 GOF Y769F)^17,23,25,26,35–37^ **(Fig. 1a)**, and in a Pol II subunit which does not directly contact Pol II active site, such as Rpb9^50^, also alter catalytic activity and fidelity, showing phenotypes seemingly identical to TL mutants, indicating that TL dynamics and function are finely balanced and could be sensitive to allosteric effects^50–53^. Determining the extent and nature of the pathways connecting the TL to the rest of the polymerase will reveal the functional networks that shape its dynamics and the potential pathways through which Pol II has evolved or may be controlled by elongation factors.

Physical and functional intramolecular interactions between amino acids define protein function and evolvability^54–57^. Genetic interactions where one mutation modulates or alters the phenotype of another are commonly referred to as epistasis. Interactions can be of different types such as suppressive, masking, or synthetic interactions. In this study, we use “epistasis” specifically to describe cases where mutant phenotypes allele-specifically depend on the identities of other amino acids. Epistasis contributes to protein evolvability by providing a physical context and an evolutionary window in which some intolerable mutations may be tolerated^58–60^. Recent studies have shown how mutations can alter protein function, allostery, and evolvability, suggesting that even conserved residues are subject to unique, context dependent epistatic constraints^61–68^. Consistent with this, distinct phenotypes for the same conserved residues have been observed in various proteins, including yeast Pol I and Pol II^66,67,69^. For example, the TL domain of yeast Pol I is incompatible with Pol II, despite sharing 70% of sequence identity, while the similarly distinct Pol III TL maintains some function^70^. It is not known generally if highly conserved TL sequences are compatible in a heterologous msRNAP context. Answering this question would reveal how tightly integrated the TL is by coevolution with its embedded context, or the extent to which its functions are self-contained due to its very high conservation.

Functional interactions between residues can be revealed by genetic interactions of double mutants^17,37,71–73^. A systematic analysis is needed to fully describe and understand the requirements for each mutant’s phenotype, and the nature of the interactions that control Pol II activity. Previous studies from our lab on a small subset of site-directed substitutions identified distinct types of Pol II double mutant interactions, including suppression, synthetic sickness and lethality (activity-additive), less-than-additive interactions (sometimes referred to as epistasis), and sign-epistasis^17,37,72^. Suppression interactions are common between LOF and GOF mutants, as expected if each mutant is individually acting in the double mutant, balancing opposing effects on activity. Similarly, synthetic lethality or sickness were common between mutants of the same class (GOF or LOF), consistent with greater defects arising from combining partial losses of TL function. However as just noted, we also observed mutant combinations where the double mutant was not sicker than the single mutants for mutants of the same class, suggesting these mutants might act at the same step. Additionally, we had observed a potential case of sign-epistasis, where a mutant’s phenotype class appeared dependent on the identity of another residue. For example, the GOF TL substitution Rpb1 F1084I was unexpectedly lethal with the LOF TL substitution Rpb1 H1085Y, contrary to the predicted mutual suppression for a GOF/LOF combination^37,72^. This suggested that F1084I required H1085 for its GOF characteristics and became a LOF mutant in the presence of H1085Y (**Fig. 1b**).

Deducing large-scale residue interaction networks is challenging. To accomplish this for Pol II, we developed experimental and analytical schemes to extend the previously established “Pol II phenotypic landscape”^17,37,72,74^ to 15,174 double and multiple mutants within the *S. cerevisiae* Pol II TL and between the TL and adjacent domains (a “Pol II Interaction landscape”). Our analyses indicate that TL function and evolution are dominated by widespread epistasis. The TL faces extensive constraints despite its high conservation, arising from residues within the TL and its interacting domains, resulting in highly similar TLs being incompatible inside yeast Pol II to a large degree. Additionally, individual mutants with similar biochemical and genetic phenotypes have quite distinct genetic interaction networks, revealing differential functional requirements for their abilities to confer altered Pol II activities. Some mutants minimally change the genetic behavior of most TL substitutions when combined, suggesting that TL conformations or interactions remain largely unchanged, with altered TL dynamics hypothesized to be the sole difference. In contrast, other alleles greatly perturb the genetic behavior of many TL substitutions, indicating these substitutions change the active site landscape in ways that would otherwise be undetectable. Together, these results suggest small changes in the Pol II active site can dramatically change its constraints and therefore, its potential evolutionary paths. Furthermore, our findings highlight potential communication networks through evolutionary coupling of Pol II residues within the active site, which may underlie allosteric communication paths to the active site for elongation factors.

## RESULTS

### A system for dissection of the Pol II active site genetic interaction landscape

We developed an experimental and analytical framework, termed the Pol II TL interaction landscape, to dissect residue interactions that shape Pol II TL function and evolution in *S. cerevisiae*. We designed and synthesized 15,174 variants representing evolutionary haplotypes, all possible Pol II TL single mutants and a subset of targeted double mutants in ten libraries (Supplemental Table 1). This approach follows our prior analysis of the TL phenotypic landscape^17^ with modifications (**Methods** and **Extended Data Fig. 1a**). Libraries were transformed, screened under selected conditions (**Methods**), and sequenced (**Fig. 1c, Extended Data Fig. 1a**). Mutant growth phenotypes are calculated based on relative allele frequency shifts compared to a control condition and normalized to the WT under identical conditions. Biological replicates indicate high reproducibility (**Extended Data Fig. 1b-c**). Individual libraries were min-max normalized^75^ to account for scaling differences between libraries (**Fig. S1a**) and the same mutants present among different libraries indicate high correlation of fitness determinations in each library (**Fig. S1b-c**).

We defined a conceptual framework for evaluating genetic interactions among TL mutations. First, we assume that independent mutant effects combine in log-additive manner. This means that predicted double mutant fitness defects should be the combination of both single mutant defects, as is standardly assumed^60,71,73,76^. Deviations from log-additive expectations represent genetic interactions. To quantify any potential deviation, we compared the observed fitness of a double mutant measured in our high-throughput experiment to its expected fitness, calculated as the addition of constituent single substitutions’ fitnesses, which are expressed as a log value. We term these differences as a “deviation score” (Deviation score = observed fitness – expected fitness; **Extended Data Fig. 2**). We consider a deviation score between -1 and 1 to be consistent with additive behavior, meaning independent contributions of single mutants to the double mutant fitness. A deviation score > 1 is taken as a positive genetic interaction (e.g., suppression), while a score < -1 indicates a negative genetic interaction (e.g., synthetic sickness or lethality).

Importantly, our phenotyping analyses determine both general mutant fitness and predict their class of in vitro biochemical defect. Our previous studies showed that phenotypic profiles from conditional growth assays, reflecting Pol II-activity dependent phenotypes in vivo, correlate with the measured biochemical activity in vitro^17,37,77^. For example, catalytically hyperactive GOF mutants have a phenotypic profile based on sensitivity to mycophenolic acid (MPA) and weak suppression of two genetic reporters (*gal10Δ56* and *lys2-128∂*), while catalytically defective LOF mutants are MPA resistant, show strong suppression of *gal10Δ56* and fail to suppress *lys2-128∂* (more fully described in Methods). Using the established correlation between growth profiles (MPA and media used to detect phenotypes of mutants on *gal10Δ56* and *lys2-128∂* backgrounds) and known/predicted in vitro catalytic effects (GOF or LOF) of 65 TL alleles^17,37,53,78^, we trained two multiple logistic regression models to distinguish Pol II mutant classes, and applied them to all viable mutants (growth fitness score > −6.5) of designed variants, classifying them into three groups, GOF (22.96%), LOF (28.11%) and unclassified (mutants not belonging to either class, 48.93%) (**Extended Data Fig. 3a**, Methods). To visually inspect the classifications, we applied t-SNE projection and k-means clustering for all measured mutants across all growth conditions. We observed separated GOF and LOF clusters consistent with logistic regression classifications (**Extended Data Fig. 3b-c**). Notably, mutants with similar defects may be further distinguished in different clusters, suggesting their similar phenotypic defects may be related to different transcription defects (**Extended Data Fig. 3c**). By classifying Pol II active site mutants as “GOF” or “LOF” based on predicted in vitro biochemical phenotypes, we establish baseline expectations for their genetic interactions. We previously observed activity-additive interactions, meaning suppression between mutants of different classes (GOF+LOF) or synthetic sickness/lethality within the same class (GOF+GOF or LOF+LOF). Together with the calculated deviation scores, we can distinguish specific epistatic interactions from activity-dependent effects. We classified an interaction as epistasis when we observed positive fitness deviation in mutants of the same activity class (GOF+GOF, LOF+LOF), where we would otherwise expect synthetic sickness or lethality if mutants were functioning independently. Conversely, we defined sign epistasis for situations where negative deviations in combinations between the classes (GOF+LOF), where suppression would be expected for independent mutants (**Fig. 1d, Extended Data Fig. 2**).

We compared interactions observed in sequencing to growth assays for 59 previously examined double mutants^17,37^, with 48 matching our observations. The other 11 were categorized as ultra-sick/lethal and therefore inaccessible to sequencing analysis (**Supplemental Table 2**). We constructed 50 additional mutants to assess interactions by patch assay and spot assay, and these aligned with our high-throughput results (**Extended Data Fig. 4**).

### Widespread incompatibility of conserved TL sequences with yeast Pol II suggests the TL is highly constrained by its coevolved environment

We previously found that identical mutations in a residue conserved between the Pol I and Pol II TLs yielded different biochemical phenotypes^70,79^. Additionally, the yeast Pol I TL was incompatible within the yeast Pol II enzyme, implying that TL function in that case depended on its enzymatic context^70,79^. To determine the scope of TL-Pol II incompatibility, we designed a library containing evolutionary TL variants from bacterial, archaeal, and eukaryotic msRNAPs and determined their compatibility in the yeast Pol II context (**Fig. 2a**). Eukaryotic TL alleles were more compatible than those from archaea or bacteria, with Pol II alleles being the most compatible (**Fig. 2b, Extended Data Fig. 5a-b**), consistent with evolutionary distance. Growth fitness in the Pol II background showed a slight negative correlation with the number of TL substitutions for most TLs but not those from bacteria (**Extended Data Fig. 5c**), likely because of the bacterial TLs being largely incompatible with Pol II. Despite this, some archaeal TLs provide yeast Pol II viability, yet several Pol II TLs were defective or lethal. These results suggest widespread coevolution between TL sequences and that the enzymatic background, beyond ultra-conserved positions, shapes TL function in individual enzymes. We have explored the nature of this incompatibility and complex intra-TL genetic interactions further in a separate study^80^.

**Figure 2.**
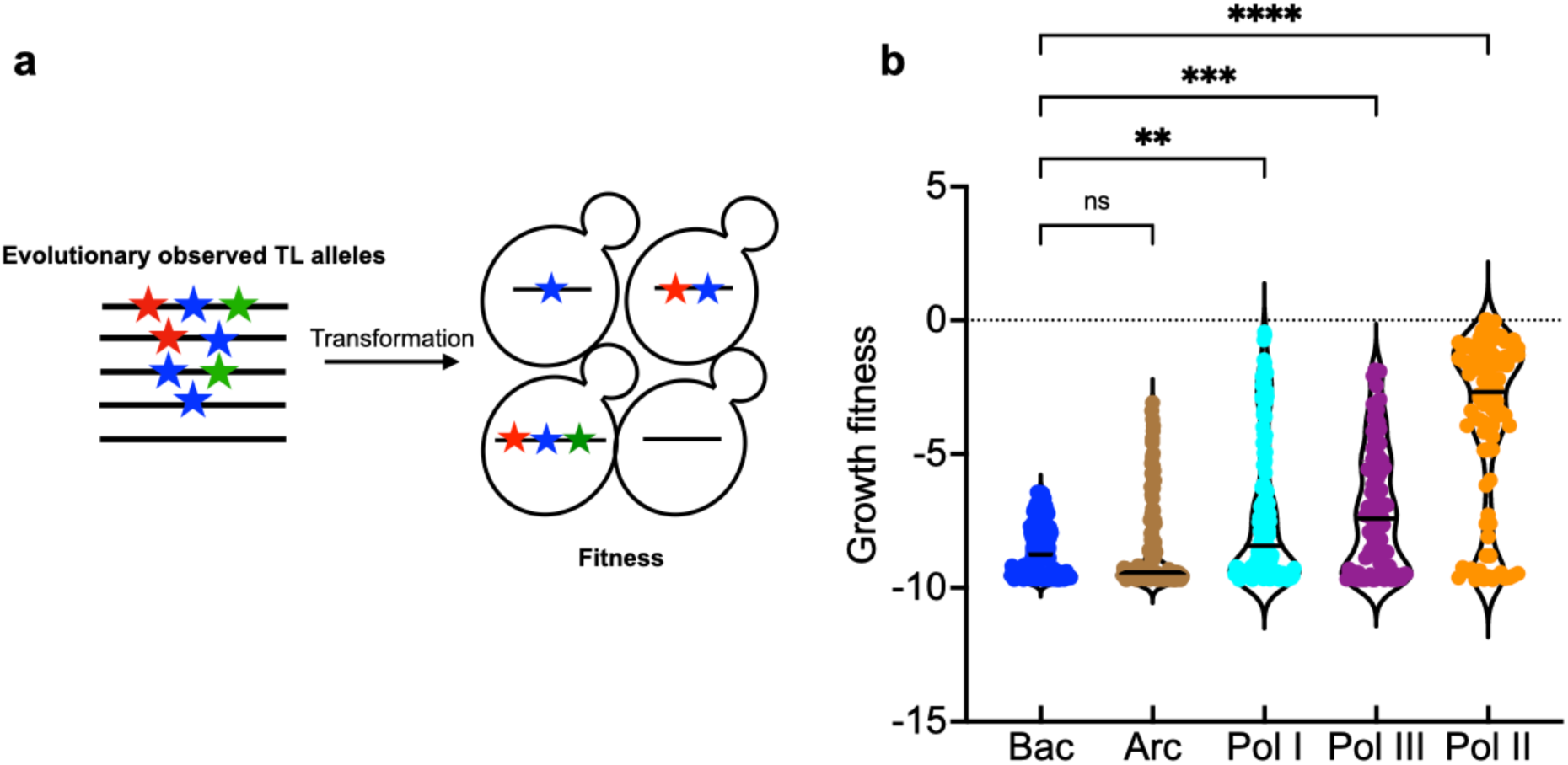
Contextual epistasis shapes TL evolution. **a.** Schematic for the TL evolutionary haplotypes library. We selected 662 TL haplotypes representing TL alleles from bacterial, archaeal and the three conserved eukaryotic msRNAPs. These TL alleles were transformed into yeast and were phenotyped under selective conditions. **b.** Fitness of evolutionarily observed TL haplotypes in the yeast Pol II background. The Pol II WT TL fitness (0) is labeled as dotted line. Kruskal-Wallis test was performed for comparison and significant levels (P < 0.05) were labeled.

### Allele-specific interactions suggest unique properties of individual mutants with similar phenotypes

To determine the potential residue interactions surrounding the TL in the yeast Pol II environment, we analyzed 12 previously studied GOF and LOF mutants (eight within the TL and four outside), each combined with >600 perturbations in the form of all single substitution TL mutants (**Fig. 3a**). These 12 mutants serve as probes for the genetic interaction space of the TL, revealing how it might be altered in allele-specific fashion by each “probe” mutation. Importantly, TL conformational dynamics and function are balanced by internal residue interactions within the TL and by TL-external interactions between the TL and TL-proximal domains. While GOF and LOF mutants in TL-adjacent domains appear similar to GOF and LOF TL mutants, their underlying mechanisms or specific residue dependencies are not known. We therefore compared interaction networks among probe mutants with similar biochemical and phenotypic defects to distinguish if changes to TL function might reflect simple alterations to TL dynamics, or additional alterations to folding trajectories or conformations.

**Figure 3.**
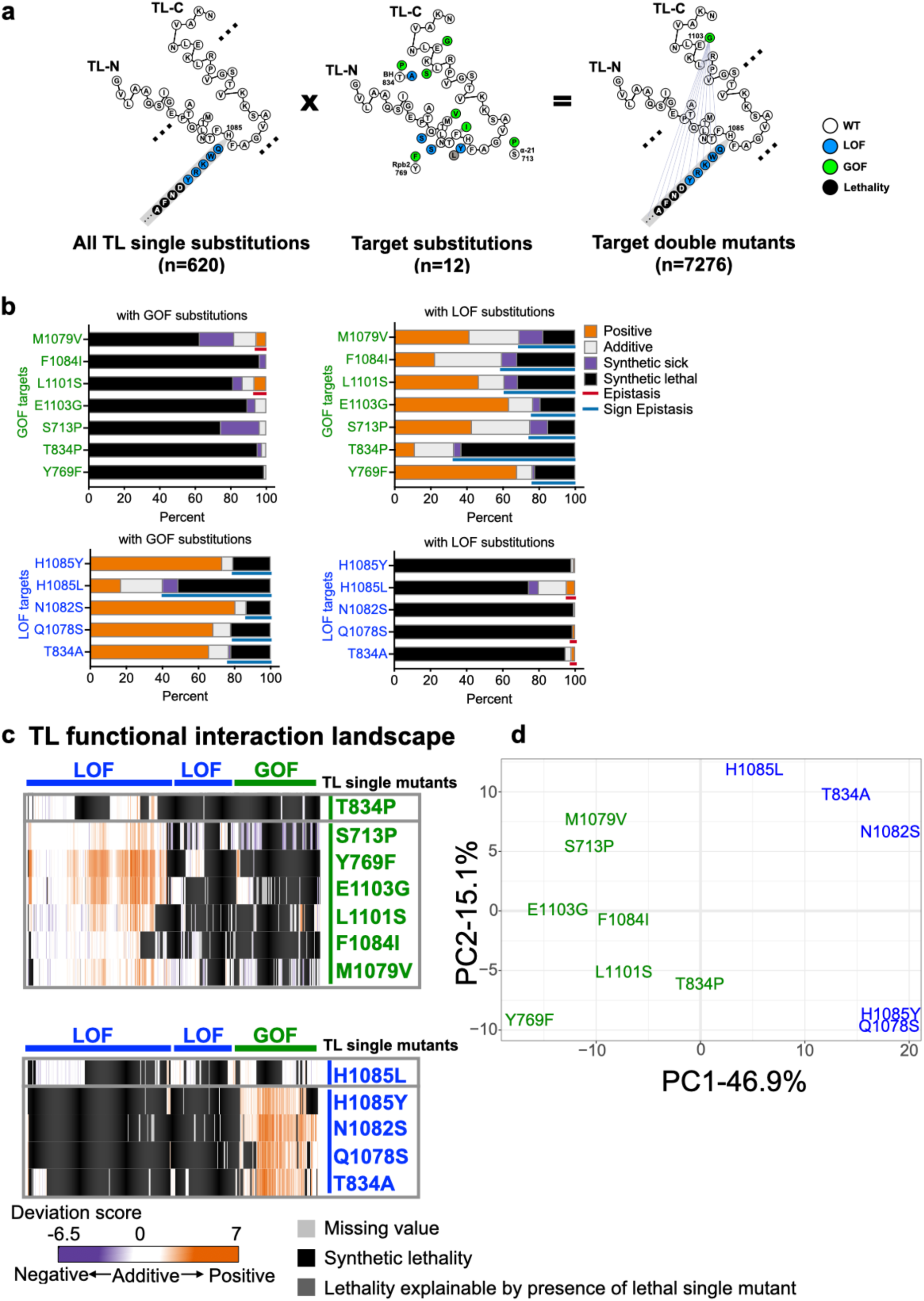
**Pol II TL interaction landscape distinguishes mutants with similar phenotypes**. **a.** Design of the targeted double mutant libraries. All possible substitutions at each TL residue (represented with a simplified format in the left panel) and twelve “probe” mutations (eight within the TL and four in TL-proximal domains) (middle panel) were combined with to generate 7280 double mutants (right panel). 7276 mutants passed the reproducibility filter and were used for interaction analyses. **b.** The percentage of functional interactions observed for each probe mutant with viable GOF or LOF TL substitutions. Epistasis and sign epistasis are labeled with colored lines. **c.** Pol II-TL functional interaction landscape with interactions represented by deviation scores. The upper panel shows interactions of GOF probe mutants in combination of viable GOF or LOF TL substitutions. The lower panel shows interactions of combinations with LOF probe mutants. **d**. Principal component analysis (PCA) of deviation scores across double mutant interactions for 12 probe mutants (see **Methods**).

We observed distinct interaction networks among probe mutants, even for those with similar apparent biochemical and growth defects (**Fig. 3b-d**), indicating that mutants differentially impact the Pol II active site, potentially reflective of different constraints and distinct properties of individual mutants, as discussed below. A simple expectation is that mutants with altered TL dynamics yet maintenance of conformational states should show additive interactions, because the effects of substitutions on function would be maintained. In contrast, substitutions altering TL conformations/folding or propagating changes across the active site, may be subject to allele-specific interactions *i.e.* prominent epistasis. We observed that most probe mutants were placed on continuum of effects based on the balance between additive (independent) interactions and epistatic (dependent or allele-specific) interactions in double mutants, which we interpret as the degree to which they propagate their effects across the active site (**Fig. 3b-c, Extended Data Fig. 6, and Fig. S2**). Probe mutants appeared separable based on the extents of their additive or epistatic interactions, where greater numbers of independent interactions (additive) would indicate fewer effects on other residues’ functions. In contrast, an increase in dependent interactions (sign epistasis) would indicate greater changes in other mutant’s characters and presumably residue function. For example, Y769F, a GOF TL-proximal mutant in Rpb2 link domain (**Fig. 1a**), uniquely suppresses some TL lethal substitutions among all GOF probe mutants (**Extended Data Fig. 7**), indicating specific functional interactions between Y769F and these substitutions. Moreover, two TL-adjacent GOF mutants, Rpb1 S713P (funnel α-helix 21) and T834P (BH) displayed largely distinct interaction networks despite similarly increased biochemical activity and genetic phenotypes as single mutants^17^ **(Fig 3c, Extended Data Fig. 8a**). S713P broadly suppressed LOF TL substitutions (96 instances), consistent with generic enhancement and preservation of TL dynamics or residue function. In contrast, T834P exhibited sparser suppression (33 instances) but much greater sign epistasis than S713P (102 versus 38 instances), especially with P1099, where all substitutions showed sign epistasis with T834P but not S713P (**Extended Data Fig. 8a**). These results suggest that T834P’s GOF function requires most TL residues to be WT, is especially reliant on proline at 1099, and that T834P alteration to the BH extensively changes the TL’s functional space, likely through folding or altered conformations.

A similar distinction was observed between two internal TL GOF mutants, Rpb1 E1103G and F1084I **(Fig. 3c, Extended Data Fig. 8b**). E1103G showed widespread suppression of LOF TL substitutions (184 instances), consistent with its proposed role in altering TL dynamics to promote TL closure^23,37^ and which allows TL mutants to maintain their effects. In contrast, F1084I showed more limited suppression (43 instances) while showing widespread synthetic lethality due to predicted sign epistasis, indicating that F1084I requires many WT residues for GOF characteristics and itself may switch to a LOF upon many additional TL perturbations.

The most striking example came from two LOF substitutions at the ultra-conserved Rpb1 H1085 **(Extended Data Fig. 8c**). This histidine, which contacts incoming NTP substrates^5,12^, is the target for the Pol II inhibitor α-amanitin^36,81^, and promotes catalysis^12,13^. Initial structural data and molecular dynamics simulations suggested that H1085 functions as a general acid for Pol II catalysis^82–85^. Our discovery that H1085L was well-tolerated^17^, and subsequent experiments from the Landick lab^86,87^, have led to their proposal that the TL histidine functions as a positional catalyst where a similarly sized leucine substitution supports catalysis with mild effects on biochemistry and growth. If H1085Y and L substitutions represent a continuum of positional catalyst activity, then their interaction networks would be expected to be similar, and differing only in magnitude, not identity or type of interactions. Contrary to this, distinct interaction patterns were observed (**Fig. 3c, Extended Data Fig. 8c**). Most GOF mutants suppressed H1085Y but not H1085L. Instead, H1085L showed synthetic lethality with most GOF mutants (putative sign epistasis). For example, almost all substitutions at E1103 showed sign epistasis with H1085L but not H1085Y (**Fig. S2b, Extended Data Fig. 8c**). The distinction between H1085L and H1085Y is further highlighted in PCA analysis (**Fig. 3d**).

### Functional and epistatic interactions within the TL hydrophobic pocket tune Pol II activity

Several allele-specific epistatic interactions were observed, with some of the most extensive occurring between A1076 substitutions and L1101S (**Fig. 4a**), suggesting tight coupling between A1076 and L1101 for Pol II function. These two hydrophobic residues, together with other hydrophobic residues in TL helices, likely stabilize the open TL conformation through hydrophobic packing of the TL helices (**Fig. 4c**). Consistent with this, another pair of adjacent residues, M1079 and G1097, also showed allele-specific interactions (**Fig. 4b-c**). Additionally, three GOF/GOF combinations between L1101S and A1076 substitutions showed WT-like behavior for the double mutants based on our regression model for mutants, indicating mutual suppression due to one of the singles switching to a different class (sign epistasis), further supporting a tight functional interaction between 1101 and 1076 (**Fig. S3a**).

**Figure 4.**
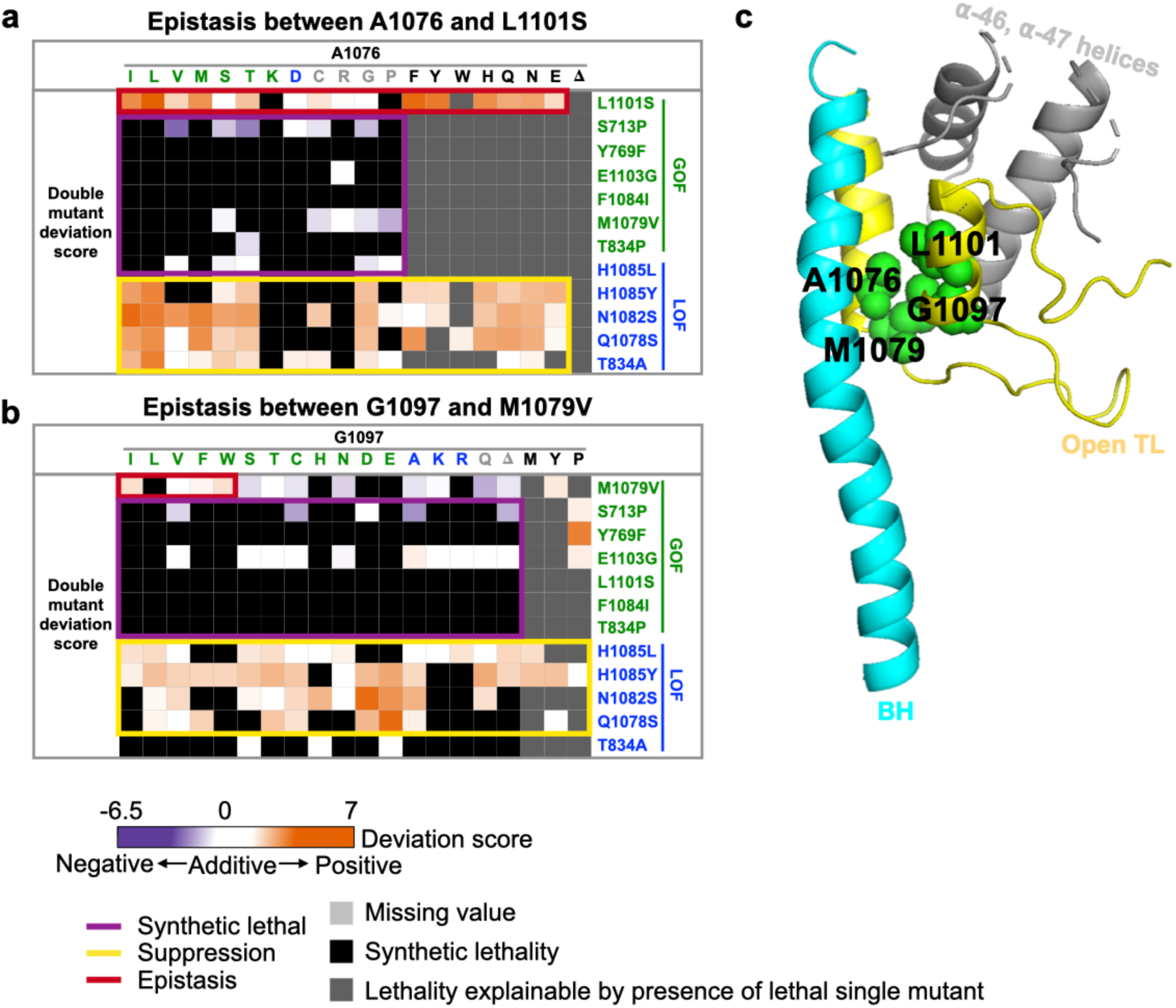
Pol II TL interaction landscape reveals functional dependency of proximal residues. a-b. Specific epistatic interactions observed between hydrophobic residues A1076 and L1101 (A), and M1079 and G1097 are shown as heatmaps (B). The x-axis of both heatmaps are 20 substitutions ordered by predicted phenotypic classes, and the color of substitution represents the phenotypic class of the substitution. GOF substitution is in green, LOF is in blue, unclassified is in gray, and lethal (fitness < −6.5) is in black. **c.** The epistatic interactions we identified between A1076 and L1101, together with M1079 and G1097 are shown on the five-helix bundle of Pol II active site (PDB: 5C4X)^42^.

We also observed allele-specific interactions for predicted lethal mutants. Our lethality threshold is more conservative than the actual lethal threshold and therefore, some classified lethal but actually slow-growing mutants have enough growth in phenotypic assays for GOF or LOF class determination. Among 21 ultra-sick/lethal TL substitutions predicted as GOF, suppression by LOF mutants was expected and observed (**Fig. S3b**). For example, lethal A1076 substitutions were suppressed by LOF probe mutants, suggesting their severe defects result from strong GOF status. However, some A1076 substitutions were also suppressed by the GOF probe mutant L1101S, indicating allele-specific mutual suppression or sign-epistasis, consistent with A1076-L1101 interactions described above (**Fig. S3c**). Similarly, ultra-sick/lethal LOF substitutions were commonly suppressed by GOF mutants, but some were suppressed by specific LOF mutants (**Fig. S3b**). For example, S1091G was suppressed by most GOF mutants and specifically by the LOF V1094D (**Fig. S3c**), suggesting allele-specific compensation. Furthermore, some lethal mutants exhibited interactions with specific GOF or LOF probe mutants. For example, F1084R, was specifically suppressed by LOF probe Q1078S and GOF probe Y769F (**Fig. S3c**). Notably, F1084, Q1078S and Y769 are in close proximity in the substrate bound, catalytic-favoring TL conformation. These types of allele-specific interactions potentially allow the TL and adjacent residues to evolve and differentiate while maintaining essential functions.

We note that strong epistasis is much more prevalent in the Pol II system than in other quantified proteins^57,88–91^ (**Fig. S3d**), likely due to the high rate of suppressive interactions stemming from Pol II mutants’ opposing effects on catalysis.

### Mapping TL-internal residue interaction networks suggests widespread epistasis within the TL

To determine TL-internal interaction networks, we selected 2-4 substitutions for each TL residue, representing diverse phenotypes (GOF, LOF, lethal, or unclassified), and combined them with the selected substitutions across all other TL positions. This curated set of 3790 double mutants captures potential interactions between any two TL residue positions (**Fig. 5a**). Observed fitness of these double mutants was compared to predictions from the additive model, revealing widespread deviation from the model (i.e. large interactions)(r^2^=0.21), much smaller than the r^2^ (about 0.65-0.75) reported in other studies^57,73,88,89,91^ (**Fig. 5b**), likely reflecting our selection criteria for mutations with diverse phenotypes. In general, mutations of the same class were additive or exacerbating, with some exceptions that reveal functional dependencies between residues (**Fig. 5c, 5e, Extended Data Fig. 9**). Additionally, the strongest deviations from expectations were between mutations of opposing phenotypic classes (GOF+LOF combinations). Here, commonly we observed suppression as predicted for mutations acting independently in 43% of GOF+LOF combinations but we also observed unexpected lethality or exacerbation (sign epistasis) for 41% of GOF+LOF combinations. Distinct patterns of sign epistasis for GOF/LOF combinations provide the ability to distinguish functional requirements for GOF alleles to be able to confer GOF phenotypes (discussed below). Interactions were distributed throughout the TL, covering every residue, supporting connectivity across the TL (**Fig. 5d**). Notably, epistasis clustered within the C-terminal TL helix and adjacent regions (The far-right panel of **Fig. 5d**), supporting functional dependencies of TL-C terminal residues on one another, consistent with their proposed function in collaboratively stabilizing the open TL.

**Figure 5.**
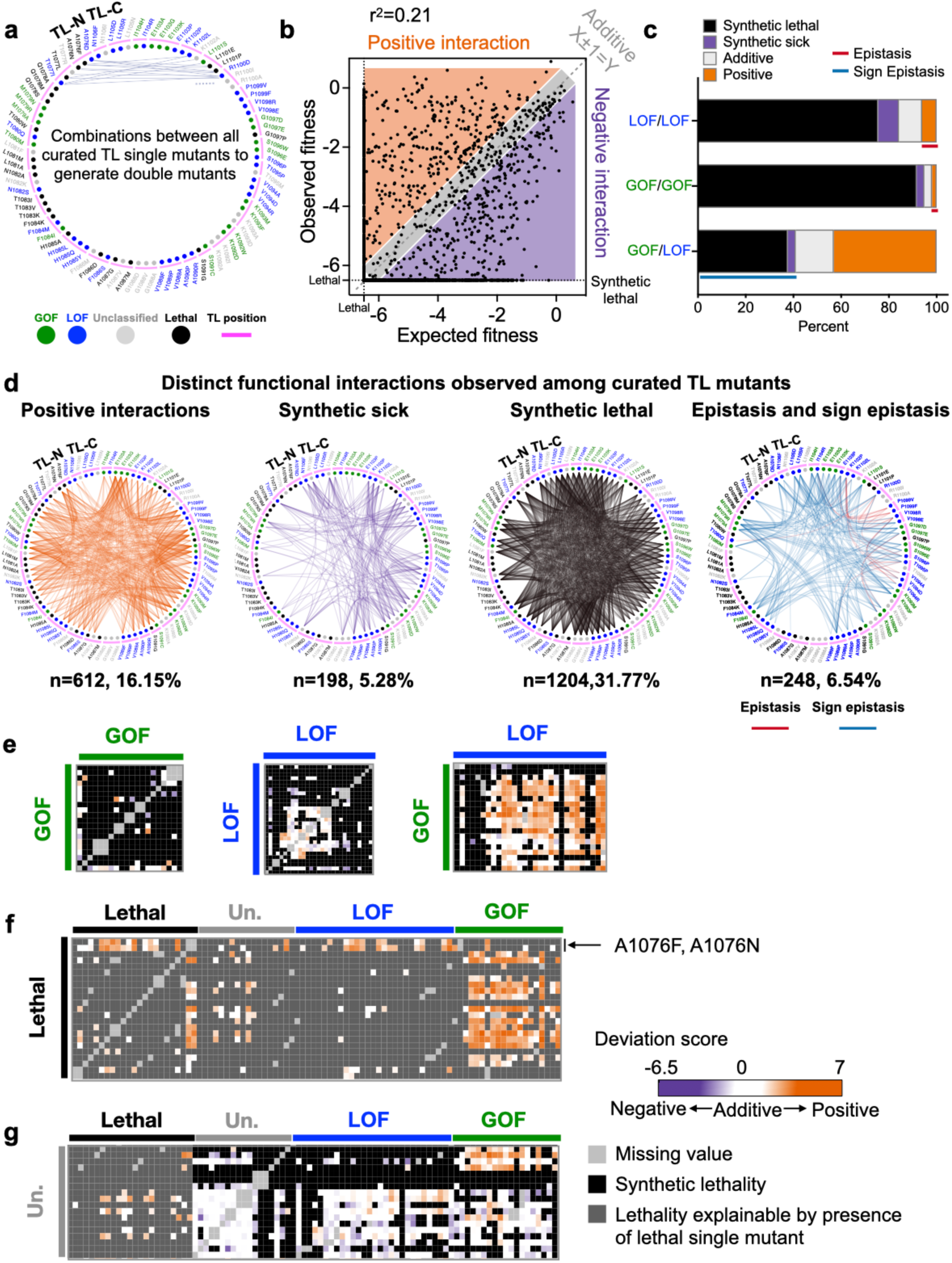
Widespread epistasis in the Pol II TL interaction landscape. **a.** Design of the pairwise double mutant library. We curated 2-4 substitutions for each TL residue (in total 90 substitutions, n(GOF) = 18, n(LOF) = 30, n(Unclassified) = 19, n(Lethal) = 23) and combined them with each other to generate double mutants. 3910 double mutants representing combinations between any two TL residues were measured and 3790 of them passed the reproducibility filter. WT TL residue positions are indicated with magenta arch. Phenotype classes of single substitutions are shown as colored circles (GOF in green, LOF in blue) while unclassified mutants are in grey and lethal mutants are in black. **b.** An xy-plot of observed double mutant growth fitness measured in our experiment (Y-axis) and expected fitness from the addition of two constituent single mutants’ fitnesses (X-axis). N (positive) = 612. N (Negative) = 1402. N (Additive) = 1776. N (Sum)=3790. Lethal threshold (−6.5) is labeled with dotted lines on X and Y axis. The additive line where X ± 1 =Y is indicated with dashed line. Simple linear regression was performed, and the best fit equation is Y = 0.52X -2.55, r^2^ = 0.21, P < 0.0001. **c.** Percent of interactions observed from each combination group. N (LOF/LOF) = 412. N (GOF/GOF) = 156. N (GOF/LOF) = 534. Epistasis and sign epistasis are indicated with colored lines. **d.** Various groups of interactions are displayed in a network format for clarity. In these networks, the TL structure is simplified to a circle, with the TL-N terminal and C-terminal of the TL connected. WT TL residues are represented by magenta arches, and selected substitutions are labeled beside these arches. The color of the label represents the phenotype of each substitution. A colored line connects two substitutions if there is interaction between them, with the line color representing the interaction type. For example, in the “Epistasis and sign epistasis” network, we selected three substitutions at K1102: K1102P, K1102L and K1102A. They show epistasis with V1094D and R, T1095P, S1096P, V1098R and P1099F and V. These interactions are labeled with red lines. The observed epistasis, indicated by red lines, accumulated at the TL-C terminal, indicating a functional dependency in this region. **e-g.** The intra-TL functional interaction heatmaps of various combinations. Double mutant deviation scores are shown in the heatmap. Annotations at the top and left indicate the curated single mutants and their predicted phenotypic classes from multiple logistic regression modeling. GOF/GOF, LOF/LOF, and GOF/LOF combinations are shown in **e**. Combinations with lethal single substitutions are in F. Combinations with unclassified mutants are in **g**.

### Genetic interactions reveal insights into lethal or unclassified individual mutants

Most lethal mutants could be suppressed by at least one predicted GOF mutant (**Fig. 5f, Extended Data Fig. 9**), suggesting their lethality likely comes from reduced activity (LOF) below a viable threshold. However, this suppressibility also suggests that individual lethal substitution activity defects are mostly close to the viable threshold. This suggests that the contributions of most individual residues represent only a portion of total TL activity, which is expected to provide a ∼1000 fold increase in catalytic activity^92–94^. Suppressibility of lethal residues by GOF alleles is also consistent with the greater probability of lethal substitutions being LOF rather than GOF. However, two lethal mutations at A1076 were suppressed by most LOF mutations or specific lethal mutants, but not GOF mutants, implying that their lethality resulted from being GOF. Additionally, unclassified single mutants generally showed limited interactions with GOF, LOF, or lethal classes, consistent with small or mostly independent effects. However, a few unclassified mutants showed suppression in combination with GOF mutants, suggesting they are either potential atypical LOF not detected by phenotypic analysis, or allele-specific interactions (**Fig. 5g, Extended Data Fig. 9**).

We reasoned that evolutionarily observed lethal substitutions would be closer to viability than those not observed and would therefore be more likely to be suppressible by Pol II GOF mutants. To determine this, we compared the suppressibility of substitutions observed in existing msRNAPs that are lethal in *S. cerevisiae* Pol II versus substitutions that are lethal but not present in extant msRNAPs by analyzing the maximum positive deviation scores from all double mutants containing individually lethal substitutions. Singly-lethal substitutions (single substitutions that are lethal on their own in yeast Pol II) identified in our msRNAP TL MSA showed higher maximum deviation scores than substitutions lethal on their own in yeast and not present in the MSA (**Fig. 6a, Extended Data Fig. 5b**). Consistently, most L1081 substitutions cannot be suppressed, whereas a L1081M, (commonly a WT residue in bacteria and archaeal RNAPs) showed the highest suppressibility (**Fig. 6b**). These results indicate that singly lethal mutants in *S. cerevisiae* Pol II observed elsewhere msRNAP evolution, on average, maintain greater functionality and/or are more easily suppressed by single additional mutants in yeast Pol II. Interestingly, most lethal substitutions in A1087 and all lethal substitutions in G1088 cannot be suppressed (**Fig. 6b**), suggesting the strict restriction at these positions, consistent with the structural observation that A1087 and G1088 are in a small pocket between the bridge and funnel helices^12,17^. The TL has been estimated to enhance catalytic activity by 500-1000 fold^92,93,95^, while yeast viability tolerates only ∼10-fold effects^36^. We conclude that lethal mutants observed as functional residues in other species are likely closer to the viability threshold, as might be predicted to result from a series of small adaptive steps that allow their function.

**Figure 6.**
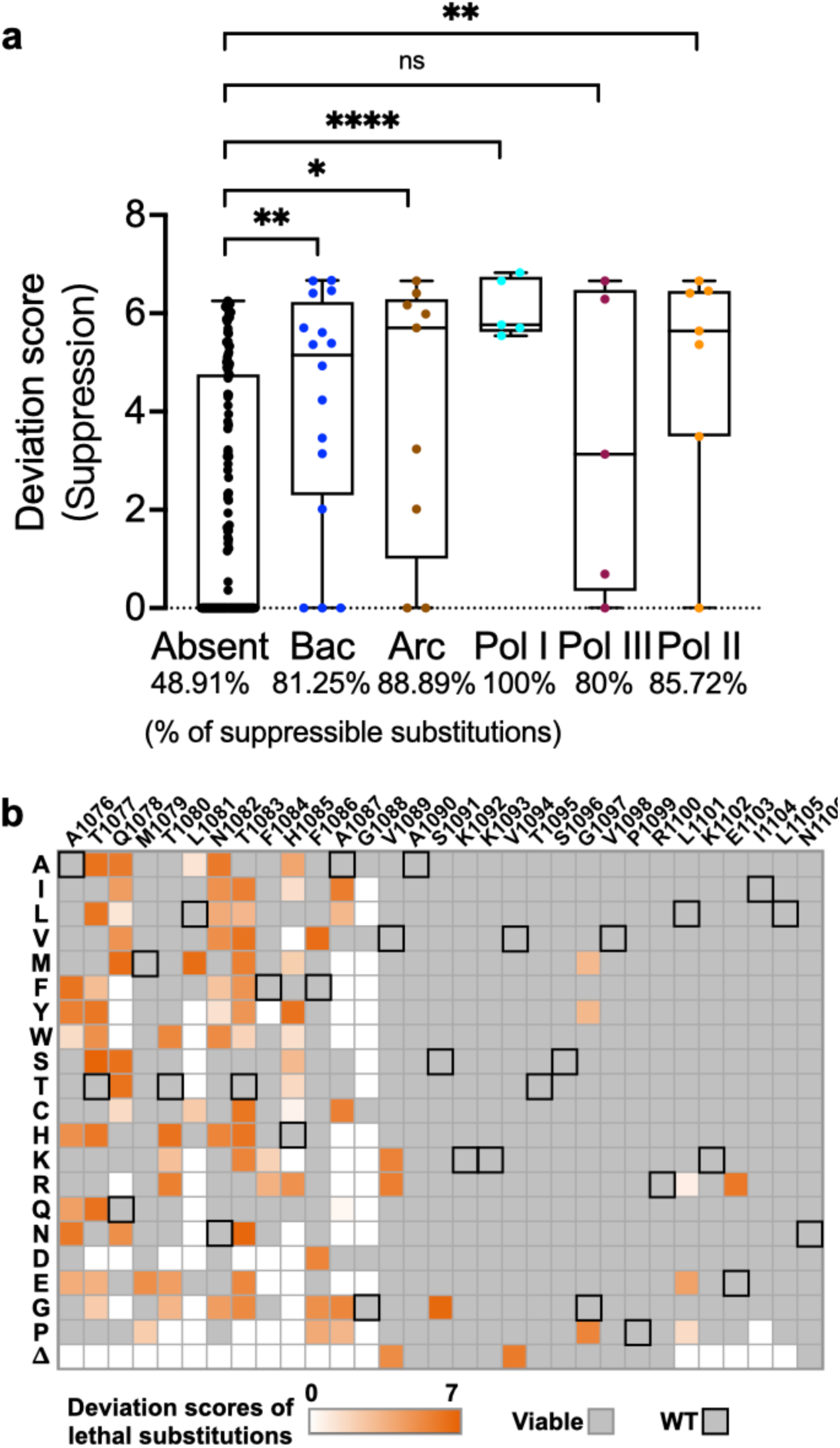
Contextual epistasis shapes TL evolution. **a.** A comparison of the maximum deviation score of each TL lethal single substitution that is present in any evolutionary TL haplotypes from bacterial, archaeal or eukaryotic Pols versus those that have not been observed in any species. The evolutionary TL haplotypes were from multiple sequence alignments (MSA). 9 substitutions were found in an MSA of 542 archaeal TL sequences that are lethal when present in yeast as a single substitution. 17 were found in an MSA of 1403 bacterial TLs, 5 were found in 749 Pol I TLs, 7 were found in 499 Pol II TLs, and 5 were found in 539 Pol III TLs. Evolutionarily observed lethal substitutions were compared to those unobserved in our TL MSA. The percentage of total suppressible lethal single mutants for each group is labeled at the bottom of the plot. Boxes are: center line, median; box limits, second and third quartiles; whiskers, maximum and minimum points. Statistical comparison was done with the Mann-Whitney test and the significant levels (P < 0.05) are shown in the figure. **b.** A heatmap displays the highest deviation scores of lethal single substitutions. The x-axis represents the wild-type residues and positions of the yeast Pol II TL, while the y-axis shows 20 substitutions. Viable substitutions are marked with gray boxes, and wild-type amino acids are indicated with gray boxes with black edge. The highest deviation scores of lethal substitutions are shown in colored boxes, ranging from white to orange, reflecting the strength of suppression.

### Co-evolution of Pol II active site residues form distinct sectors within specific domains and extend to the surfaces of Rpb1 and Rpb2

Our analyses suggest that even a highly conserved domain like the Pol II TL can be tightly integrated through coevolution of less conserved residues and that substitution tolerance is shaped by widespread epistasis. Changes in residue identities alter interaction networks, shaping the Pol II active site across evolution. To identify the coevolving residue networks in Pol II active site we employed statistical coupling analysis (SCA) for residues within Rpb1 and Rpb2, the two largest subunits of Pol II, which include the entire active site. SCA is well suited for identifying coupled residue sectors, representing putative residue communication networks^96–98^. From 283 linked Rpb1/Rpb2 sequences, we identified 53 coevolving sectors (**Fig. S4**). Residues within the active site form four major sectors (**Fig. 7a-e**). First, sector 37 includes many Rpb2 link domain residues and most TL residues, including those at the NIR and some at the TL C-terminal (**Fig. 7b**). This sector tightly surrounds the substrate binding pocket (**Extended Data Fig. 10a**), befitting critical roles in catalysis. Second, sector 15 consists of four BH residues, most residues in switch 1 (part of the clamp domain), and many cleft residues, including the ⍺46-47 loop, which accommodates incoming DNA^8,13^ (**Fig. 7c**). Residues in this sector directly interact with Pol II subunits Rpb5 and Rpb6 (**Extended Data Fig. 10b**), consistent with Rpb5’s role in interacting with the incoming DNA^99^. While the ⍺46-47 loop forms a five-helix bundle with two TL helices and the BH helix^42^ (**Fig. 4c**), coupling analysis indicates it is primarily evolutionarily coupled with cleft residues instead of the TL. This suggests that the ⍺46-47 loop may influence TL function by constraining the space for TL movement, rather than directly interacting through specific residue identities. Third, sector 1 involves active site residues surrounding the Mg^2+^ at the catalytic site and most F-loop residues (**Fig. 7d**). This sector forms a potential communication path that wraps around the TL and BH, connects to the BH through two BH residues, and extends to Rpb9, itself a subunit that forms important inputs to the TL and is required for transcription fidelity^50,100,101^. This pathway likely mediates communication between Rpb9 and the Pol II active site (**Extended Data Fig. 10c**). Fourth, sector 2 contains most BH residues are coupled with the Rpb2 Fork and hybrid binding domain, forming a network linking the BH to Rpb2 residues (**Fig. 7e**). This network connects to Rpb5, which interacts with the downstream DNA, Rpb6, a shared subunit in three eukaryotic msRNAPs, Rpb9, and Rpb3 and Rpb10, which function in Pol II assembly^102^ (**Extended Data Fig. 10d**), representing a residue communication path potentially tied to various transcription activities.

**Figure 7.**
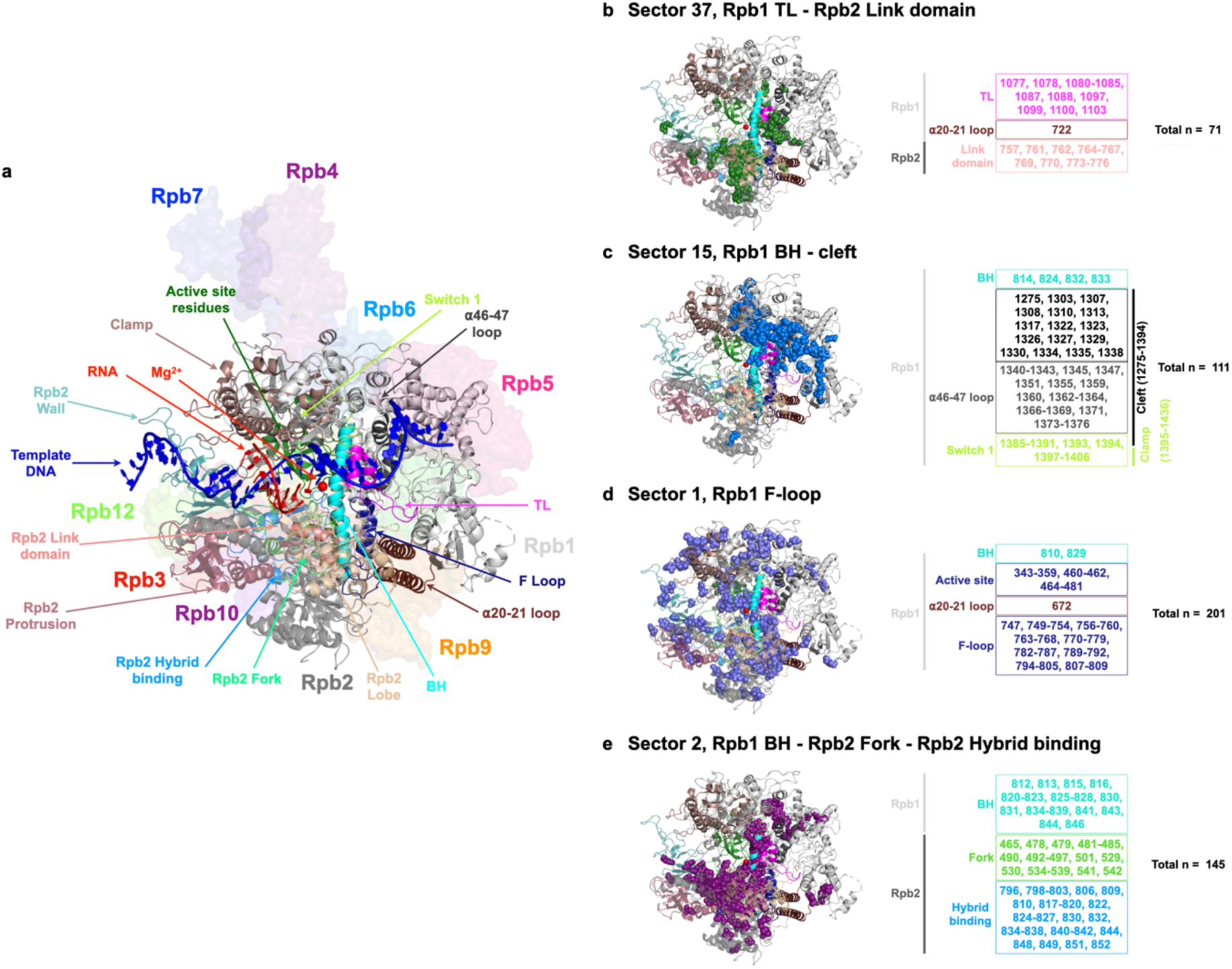
Ultra-conserved TL co-evolves with Pol II residues through diverse pathways. **a.** Structure representations of Pol II active site. Pol II subunits are showed with surface in different colors. The active site domains are in cartoon format. Rpb1 active site residues 346-375 and 436-508. Rpb1 TL, 1076-1106. Rpb1 BH, 810-846. Rpb1 ⍺20-21 loop, 672-737. Rpb1 F loop, 747-810. Rpb1 ⍺46-47 loop, 1339-1378. Rpb1 switch 1, 1385-1407. Rpb1 clamp, 1-95, 235-346, and 1395-1436. Rpb2 link domain, 757-776. Rpb2 Fork, 465-547. Rpb2 hybrid binding, 750-852. Rpb2 wall, 852-973. Rpb2 protrusion, 44-218. Rpb2 lobe, 218-405. **b-e**. Four major coevolution sectors of Pol II active site residues were identified through statistical coupling analysis. The residues within each active site domain and the total number of residues in each sector are labeled. These sectors are mapped on the yeast Pol II Rpb1/Rpb2 structure (PDB: 5C4X)^42^.

Notably, coupling is not restricted to residues that are within or near the active site. Some sectors extend from the active site to the surface of Rpb1/Rpb2 (**Fig. S5a-f**), suggesting that active site function, potentially through the TL, could be modulated by allosteric interactions through the binding of transcription factors. Residues at interfaces with elongation factors can be coupled with those factors yet some residues at interfaces can be coupled with residues distal to the interface, suggesting potential pathways for allosteric interactions. Examples of these are shown for TFIIS (**Fig. S6**), LEO1 (Leo1), ELOF1 (Elf1), and SUPT5H (Spt5) (**Fig. S7**). Together, our findings highlight potential communication networks through evolutionary coupling that may underlie regulation paths from Pol II surface to active site.

## DISCUSSION

How individual mutants alter a protein’s function is not necessarily straightforward at the mechanistic level. Amino acid substitutions both remove functionality of the WT residue but replace that functionality with something different. By altering the local environment within a protein or potentially propagating effects to distant locations, each substitution potentially can be quite different. These differences may not be apparent as phenotypic outputs, and phenotypic assays may not have granularity to distinguish different biophysical behaviors if they result in similar outputs. For Pol II mutants, even high-resolution phenotypic analyses, such as gene expression profiling or genetic interaction profiling between Pol II mutants and deletions in other yeast genes^53^, suggest that LOF and GOF mutants represent a continuum of defects that match enzymatic activity in vitro. Therefore, these profiles also appear dependent on the output of Pol II activity defects and can’t distinguish potential differences in underlying mechanism.

Through systematic detection of genetic interactions within the Pol II active site, we have identified functional relationships between amino acids across the TL and between TL substitutions and others. In the absence of double mutant epistasis analyses it would not be possible to differentiate similar alleles from one another. L1101S and E1103G, for example, are two GOF alleles very close to each other in Pol II structure and confer similar phenotypic landscapes across various growth conditions. Here, we find that their distinct interactions support that substitutions at 1101 and 1103 target distinct residue networks (**Fig. 3c, 4a, Extended Data Fig. 6, Fig. S2**). L1101 functions in the TL hydrophobic pocket while E1103 interacts and co-evolves with a number of TL external residues that together support interactions that maintain the open TL conformation. We also observed connections between TL C-terminal residues that suggest a limit to how disruptions to structure there can alter Pol II activity (**Fig. 5d**). Helix-disrupting LOF proline substitutions in at least two TL positions showed epistasis with multiple substitutions in the back of the TL (1094-1098), suggesting that their functions require TL C-terminal helix structure and in the absence of that structure (proline disruption) effects are no longer additive.

The strongest epistatic interactions were observed between two pairs of hydrophobic residues, A1076 and L1101, and M1079 and G1097 (**Fig. 4**), suggesting their interdependence in maintaining TL open conformation stability^42^. Considering the TL’s multiple active site roles, the observed interaction networks may also contribute to other NAC steps or off pathway steps (*eg.* pausing or backtracking). Structural studies have proposed that these hydrophobic residues form a pocket stabilizing the TL by bundling two TL helices with the BH and ⍺46-47 loop in a five-helix bundle^28,42^. However, our coupling analysis suggests that ⍺46-47 loop is evolutionarily coupled with cleft residues rather than the TL, indicating the ⍺46-47 loop may indirectly impact TL function by limiting the space available for TL movement, rather than through direct interactions with TL residues (**Fig. 7c**).

Elongation factors bind Pol II and alter its activity, but the mechanisms by which they do so are not known^103,104^. We observed a high level of genetic interactions between residues outside the TL and residues within it, including allele-specific reshaping of TL mutant space upon single substitution outside the TL (**Fig. 3**). The fact that minor mutational changes outside the TL can apparently functionally perturb the TL would be consistent with the idea that minor alterations to Pol II structure upon elongation factor binding could easily propagate into the active site via the TL or the BH. As an example, human Rtf1 has been observed to project a domain into the Pol II structure adjacent to the BH (in yeast, this region is occupied instead by Rpb2^105^). These contacts have been proposed to alter Pol II activity. We would propose that the paths for such alteration to activity may follow the coupling sectors we have observed by SCA (**Fig. 7**).

How different individual substitutions are under the surface is critical for understanding plasticity in protein mechanisms and how they might be altered by evolutionary change. A key open question in nucleic acid polymerase mechanisms is the paths for protons in the reaction (for example, deprotonation of the synthesized strand 3′-OH and protonation of pyrophosphate leaving group, for example^82,83,85,87,106–108^). For msRNAPs, the association with the incoming NTP by a universally conserved histidine led to the proposal that this residue might donate a proton during the reaction ^12,84,106^. Some substitutions at this position can provide minimal essential function (e.g. tyrosine, arginine), while others are only moderately defective (glutamine). Surprisingly, we found that H1085L was very-well tolerated for growth ^17^ and the Landick lab has proposed this substitution supports catalysis through positional but not chemical effects^86,87^. Our studies here were quite surprising in that they indicated that L1085 Pol II has unique behavior when perturbed by all possible TL substitutions and is entirely distinct from H1085Y (where we have direct observations of all possible intra-TL doubles) or H1085A or H1085Q (curated doubles) (**Extended Data Fig. 8c, Fig. S8**). These residue specific behaviors suggest that each substitution may have different properties, and compatibility with function may not necessarily represent similar function under the surface.

Evolutionary change over time can alter protein function but it can also alter protein functional plasticity. Recent work from the Thornton lab elegantly demonstrates that phenotypes of substitutions to residues conserved over hundreds of millions of years can change over evolutionary time and can do so unpredictably and transiently during evolution^62^. msRNAPs have structures and functions conserved over billions of years, and deep within their active sites is a mobile domain, the TL, that has large functional constraints on its sequence. The TL sequence must be able to fold into multiple states and maintain recognition of the same substrates across evolutionary space and is shows high identity even between distantly related species. Here we show that the TL, and likely the entire Pol II active site, exhibits a great amount of plasticity through non-conserved positions that are essential for compatibility of the TL and surrounding domains. A limitation of our study is its reliance primarily on genetic and statistical coupling analysis. Future biochemical and biophysical studies will give mechanistic insight into the activity defects underlying the functional connections we observe, for example connecting kinetic properties or structural states to specific mutants. Nonetheless, our results illustrating widespread epistasis and allele-specific effects of single and double mutants predict that comparative analyses among Pol I, II, and III will reveal shared but also enzyme-specific mechanisms due to higher order epistasis shaping the functions of conserved residues.

## ACKNOWLEDGMENTS

We thank Dr. Anne-Ruxandra Carvunis (U. Pittsburgh) and Dr. Steve Lockless (Texas A&M) for discussions and advice. We thank Zhizhen Wang and Muyao Lin from the Pitt Statistical Consulting Center for their advice on checking the reproducibility of our data. We acknowledge funding from NIH R01GM097260 for initiation of this project and NIH R35GM144116 for this work. This research was supported in part by the University of Pittsburgh Center for Research Computing, RRID:SCR_022735, through the resources provided. Specifically, this work used the HTC cluster, which is supported by NIH award number S10OD028483.

## AUTHOR CONTRIBUTIONS

BD performed sequencing library construction, data analysis, and made figures, drafted and revised the manuscript. CQ designed the mutant libraries, performed screening experiments, and contributed to revising the manuscript. SHZ provided allele counts from raw sequencing data. CDK conceived the project, guided analyses and interpretation of data, provided funding, and revised the manuscript.

## METHODS

### Design and Synthesis of TL mutant libraries

We updated and extended the fitness dataset of Qiu *et al*^17^. sing a similar methodology, but with adjusted conditions and a second-generation mutant library strategy to generate a complete Pol II TL mutation-phenotype map and examine genetic interactions. Mutants were constructed by oligo synthesis (Agilent) and screened for phenotypes previously established as informative for Pol II mutant biochemical defects. Programmed oligonucleotide library pools included all 620 single TL residue substitutions and deletions for Rpb1 amino acids 1076-1106 (Library 1), 3914 pairwise double substitutions (Library 2), 4800 targeted double substitutions (Library 6), and 3373 multiple substitutions (Library 3-5), along with the WT *S. cerevisiae* Pol II TL allele at a level of ∼15% of the total variants, enabling precise quantification (see Supplemental Table 1, 3). Each synthesized region contained a mutated or WT Pol II TL sequence and two flanking regions at the 5′ and 3’ ends of the TL-encoding sequence. These flanking regions also contained designed “PCR handle” (20bp) sequences, allowing distinct subsets of oligos to be amplified from synthesized pools using selected primers for PCR, and additional flanking WT Pol II sequences allow for further extension of homology arms by PCR “sewing” (Details are in Supplemental Method 3).

### Introduction of Libraries into yeast

Synthesized mutant pools were transformed into yeast (CKY283) along with an *RPB1*-encoding plasmid where the TL-encoding sequence was replaced with an *Mlu*I restriction site for linearization as described in Qiu et al^17^. This strategy allows construction of *rpb1* mutant libraries by gap repair between library fragments and the linearized vector. Briefly, the synthesized oligo pools were amplified by limited cycles of emulsion PCR to limit template switching. Extension of flanking homology arms of ∼200 bp were added by PCR sewing. Amplified TL sequences with extended flanking regions were co-transformed with linearized pRS315-derived *CEN LEU2* plasmid (pCK892) into CKY283, allowing gap repair via homologous flanking regions. In our experience phenotypes of integrated alleles and single copy plasmids are qualitatively very similar but this has not been systematically compared quantitatively. To detect potential residue-residue interactions between the TL and TL-proximal domains including the Rpb1 Bridge Helix (BH), Funnel Helix alpha-21 and Rpb2, the Pol II TL single mutant pool (Library 1, 620 mutant alleles and 111 WT alleles) was co-transformed individually with gapped plasmids encoding an additional *rpb1* allele T834P (Library 7), T834A (Library 8), or Funnel Helix alpha-21 S713P (Library 10)) into CKY283 respectively, or with the gapped WT *RPB1* plasmid into a strain with the genomic mutation, *rpb2* Y769F (Library 9). These co-transformations created double mutants between the TL and TL-proximal mutants. The WT allele in single mutant pool represented the single probe mutant due to substitutions outside the TL on the plasmid or in the strain background. To distinguish between a fully WT TL and a WT TL representing the TL of a mutant allele elsewhere, a WT Pol II TL allele with a silent mutant at T1083 (WT codon ACC was replaced with ACT) was co-transformed with plasmid containing gapped WT *RPB1* in a WT strain in parallel. 15% of the transformants with silent mutation were mixed with transformants of double mutants. The silent mutation allowed us to distinguish the WT and the single mutants.

### Phenotyping libraries with different conditions

Each transformation was done in three biological replicates. After transformation, Leu^+^ colonies were collected from **SC-Leu** plates and replated on **SC-Leu+5FOA** to select for cells having lost the *RPB1 URA3* plasmid and to measure growth defects. 5-FOA-resistant colonies were then scraped and replated on different conditions to assess functional differences between WT, GOF, and LOF mutants^17,37^. **SC-Leu + 20 μg/L MPA** (Fisher Scientific) evaluates transcription start site (TSS) shifting. MPA depletes GTP levels in yeast. To generate GTP, *IMD2* expression is required, but its promoter has multiple transcription start sites (TSS)^109,110^. Transcripts from the downstream TSS express functional *IMD2* upon MPA treatment. WT and LOF mutants are resistant to MPA due to their ability to use downstream TSSs for *IMD2* expression, while GOF mutants are sensitive as they can only use upstream TSS sand fail to express functional *IMD2*^53,77^. **SC-Leu + 15 mM Mn** (Sigma): Mn^2+^ compromises transcription fidelity^111^ and likely also increases Pol II catalysis; while WT and LOF mutants are resistant, GOF mutants are sensitive either due to decreased fidelity or exacerbated catalytic increase. **YPRaf** and **YPRafGal**: the *gal10Δ56* allele compromised *GAL10* 3’-end formation and polyadenylation, leading to transcription read-through that inhibits the downstream *GAL7* gene ^112,113^. The transcription readthrough in WT *gal10Δ56* cells leads to accumulation of a toxic galactose metabolite because of defects in *GAL7* expression. Some GOF and most LOF mutants show varying galactose resistance, likely through increasing termination in the *GAL10-GAL7* intergenic region or otherwise reducing read-through interference with *GAL7*. YPRaf serves as a control for YPRafGal, to demonstrate that the defect on YPRafGal is galactose toxicity. **SC-Lys** tests lysine auxotrophy. The *lys2-122∂* allele in the strain background has a *Ty1* transposable element insertion, which prevents normal expression of *LYS2*^114^. WT and LOF mutants are Lys^-^, while GOF mutants utilize a normally silent promoter within the *Ty1* ∂ element insertion, becoming Lys^+^. **SC-Leu + 3% formamide** (JT Baker), formamide destabilizes protein hydrogen bonds^115^, with most mutants showing resistance except for slight sensitivity in LOF mutants. Details of cell numbers plated on each plate and screening time of each plate are in Supplemental Table 5. Details of the high efficiency transformation protocol are in Supplemental Method 1.

### Generation of libraries for quantification by amplicon sequencing

Genomic DNA of each screened library was extracted using the Yeastar genomic DNA kit according to manufacturer’s instructions (Zymo Research). To ensure adequate DNA for sequencing, the TL regions of all libraries were amplified with PCR cycles that were verified to be in the linear range by qPCR to minimize disturbance of allele distributions, and under emulsion PCR conditions (EURx Micellula DNA Emulsion & Purification (ePCR) PCR kit) to limit template switching. Details are in Supplemental Method 2 and 3. To multiplex samples, we employed a dual indexing strategy wherein 10 initial barcodes for differentiating 10 mutant libraries were added during the initial amplification using 10 pairs of custom primers. In a second amplification, 28 primers containing 28 NEB indices were used to add a second index for distinguishing conditions and replicates (NEBNext Multiplex Oligos for Illumina) (see Supplemental Table 4). As a result, a sample-specific barcodes were present for each set of variants. The indexed, pooled samples were sequenced by single end sequencing on an Illumina Next-Seq (150nt reads). On average, over 11 million reads were obtained for individual samples with high reproducibility from two rounds of sequencing.

### Data cleaning and fitness calculation and normalization

Reads of mutants were sorted into appropriate libraries and conditions by detecting particular indices after sequencing. Read counts were estimated by a codon-based alignment algorithm to distinguish reads that exactly matched designated codons of mutants ^116^. To clean the data, mutant reads with coefficients of variation greater than 0.5 in the control condition (SC-Leu) were excluded from the analysis. The mutant read count was increased by 1 to calculate the allele frequency under different conditions. To measure and compare the phenotypes of all mutants, mutant phenotypic score (fitness) was calculated by allele frequency change of a mutant under selective conditions relative to the unselective condition comparing to the frequency change of WT. The formula for calculating fitness is shown below.

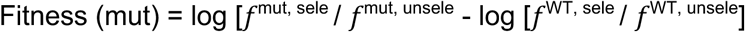

We applied min-max normalization to bring the median growth fitness of mutants measured at ten libraries to the same level for direct comparison (formula is shown below). In each library, we divided mutants into several groups based on their allele counts on the control condition. Mutants with read count differences of less than 10 are present in one group. The WT growth fitness was set as the maximum value and the minimum fitness in each group was the minimum. Min-max normalization was used to equalize the growth fitness into the same range between various groups inside each library. Additionally, we utilized min-max normalization to level the mutant fitness across all ten libraries with WT fitness as Max and minimal fitness in each library as the minimum. As a result, mutant growth fitness was scaled to one range and could be used to determine genetic interactions.

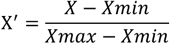

### Determination of functional interactions

The genetic interactions between single substitutions were determined by comparing the multiple-substitution mutant normalized median fitness to the log additive of the single substitution normalized median fitness. The simplified formula is as follows:

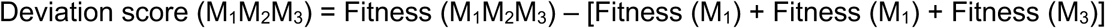

1. 1 < Deviation score < 1, the interaction among the constituent single mutants is additive and mutants are acting independently.
2. Deviation score ≥ 1, the interaction is non-additive and is positive, including suppression and epistatic interactions.
3. Deviation score ≤ -1. the interaction is non-additive and is negative, including synthetic sick, synthetic lethal, and sign epistasis interactions.

Any mutation with fitness smaller than the lethal threshold (−6.50) was classified as an ultra-sick/ lethal mutant and its fitness was normalized to −6.50 for calculation of the deviation score.

Synthetic sickness and synthetic lethality were distinguished by whether a double mutant is viable or lethal (fitness is greater than or equals to the lethal threshold −6.5) when two constituent mutations are viable. Synthetic lethality can be further classified into two types. First, additive-synthetic lethality was determined when the expected double mutant fitness calculated by additive model was lethal (expected fitness = -6.5) and the observed double mutant fitness was also lethal (fitness = -6.5) (in this case the deviation score = 0). Second, the beyond-additive synthetic lethality was determined when the expected double mutant was viable (expected fitness > -6.5) while the observed double mutant fitness was lethal (fitness = -6.5) (in this case the deviation score < 0). To separate these two situations in our figures, we labeled additive synthetic lethality as black and beyond-additive synthetic lethality as purple. Details of formulas are in Supplemental Method 4.

### Mutant classification using two multiple logistic regression models

We trained two multiple logistic regression models to distinguish GOF and LOF mutants using phenotypic fitness from SC-Leu+MPA, SC-Lys, and YPRafGal conditions of 65 single mutants. This dataset included 20 mutants with transcription rates measured by biochemical experiments (13 GOF and seven LOF)^17,37,53^, and 45 additional mutants that were hierarchical clustered with verified mutants (12 GOF, 26 LOF, 1 WT, and 6 that were not GOF or LOF mutants)^17^. Both models incorporated intercept, main effects, and two-way interactions were involved in defining both models, with a cutoff threshold of 0.75 applied to classify mutants in both of the GOF and LOF models.

Model for predicting the probability of a mutant being a GOF:

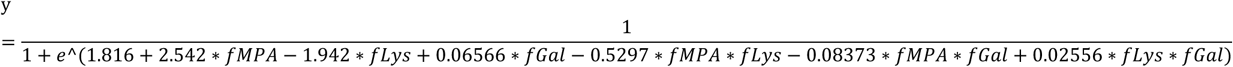

Model for predicting the probability of a mutant being LOF:

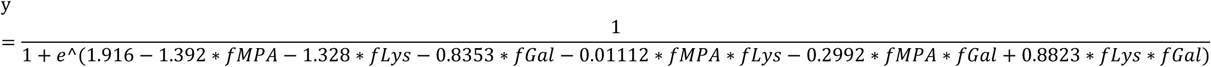

Both models showed accuracy, with the area under ROC close to one **(Extended Data Fig. 2A**). The details are provided in Supplemental Table 6.

### Principal component analysis (PCA)

Deviation scores of curated and probe double mutants were analyzed in PCA. The scripts used R language v4.0.3 (https://www.R-project.org/) with R packages tidyverse v1.3.1 (https://www.tidyverse.org), prompt (stats v3.6.2 (https://www.rdocumentation.org/packages/stats/versions/3.6.2/topics/prcomp)), ggplot2 v3.3.3 (https://ggplot2.tidyverse.org), dplyr v1.0.6 (https://dplyr.tidyverse.org), and missMDA v1.18 (https://dplyr.tidyverse.org).

### t-SNE projection

Allele frequencies for all mutants in nine conditions with three replicates were analyzed by t-SNE (Perplexity = 50) or k-means (clusters =20). Thirteen clusters with ultra-sick to lethal mutants as majority were eliminated. The remaining mutants were analyzed again with t-SNE (Perplexity = 100) and k-means (cluster =10). The scripts utilized R language v4.0.3 (https://www.R-project.org/), along with R packages Rtsne v0.15 (https://github.com/jkrijthe/Rtsne), ggplot2 v3.3.3 (https://ggplot2.tidyverse.org), k-means (stats v3.6.2 (https://www.rdocumentation.org/packages/stats/versions/3.6.2/topics/kmeans)).

### Statistical coupling analysis

5787 eukaryotic homologous sequences of yeast Rpb1 from a published multiple sequence alignment (MSA) and 4000 Rpb2 sequences from NCBI nr database were cleaned and used in the statistical coupling analysis^87^. Sequence identity was reducing to 90% with T-coffee package v12.00.7fb08c2^117^. Pol I, II, and III sequences were separated based on a phylogenetic tree constructed with FastTree 2^118^. Finally, 283 matched Pol II Rpb1 and Rpb2 sequences, with each matched pair sharing the same species name, were connected with 10 Ns and aligned with T-coffee. This newly generated MSA was used for statistical coupling analysis with the python-based package pySCA v6.1^97^. The scripts were adapted from https://github.com/ranganathanlab/pySCA.

## Supplemental Tables

Supplemental Table 1. Library summary

Supplemental Table 2. Double mutant interactions observed in high throughput system and growth assays in published papers

Supplemental Table 3. Strains and plasmids Supplemental Table 4. Primers Supplemental Table 5. Phenotyping details

Supplemental Table 6. MLR_models_summary

Supplemental Table 7. Kruskal-Wallis tests

## Supplemental methods

Supplemental Method 1. High efficiency large scale chemical yeast transformation protocol.

Supplemental Method 2. Emulsion PCR set up with EURx Micellula DNA Emulsion & Purification (ePCR) PCR kit.

Supplemental Method 3. Amplification/transformation/screening of mutant libraries and sequencing pool preparation.

Supplemental Method 4. Formulas of calculating functional interactions.

## DATA AVAILIABILITY

Raw sequencing data has been deposited on the NCBI SRA (Sequence Read Archive) database under BioProject PRJNA948661. Processed mutant count, fitness and processing codes are available through GitHub (https://github.com/Kaplan-Lab-Pitt/TLs_Screening.git). Strains and plasmids will be provided upon request. Source data are provided with this paper.

## CODE AVAILIABILITY

The codes for calculating deviation scores, PCA, t-SNE projection, statistical coupling analysis, and generating figures are available in GitHub (https://github.com/Kaplan-Lab-Pitt/TLs_Screening.git).

---

**Extended Data Fig.1.**
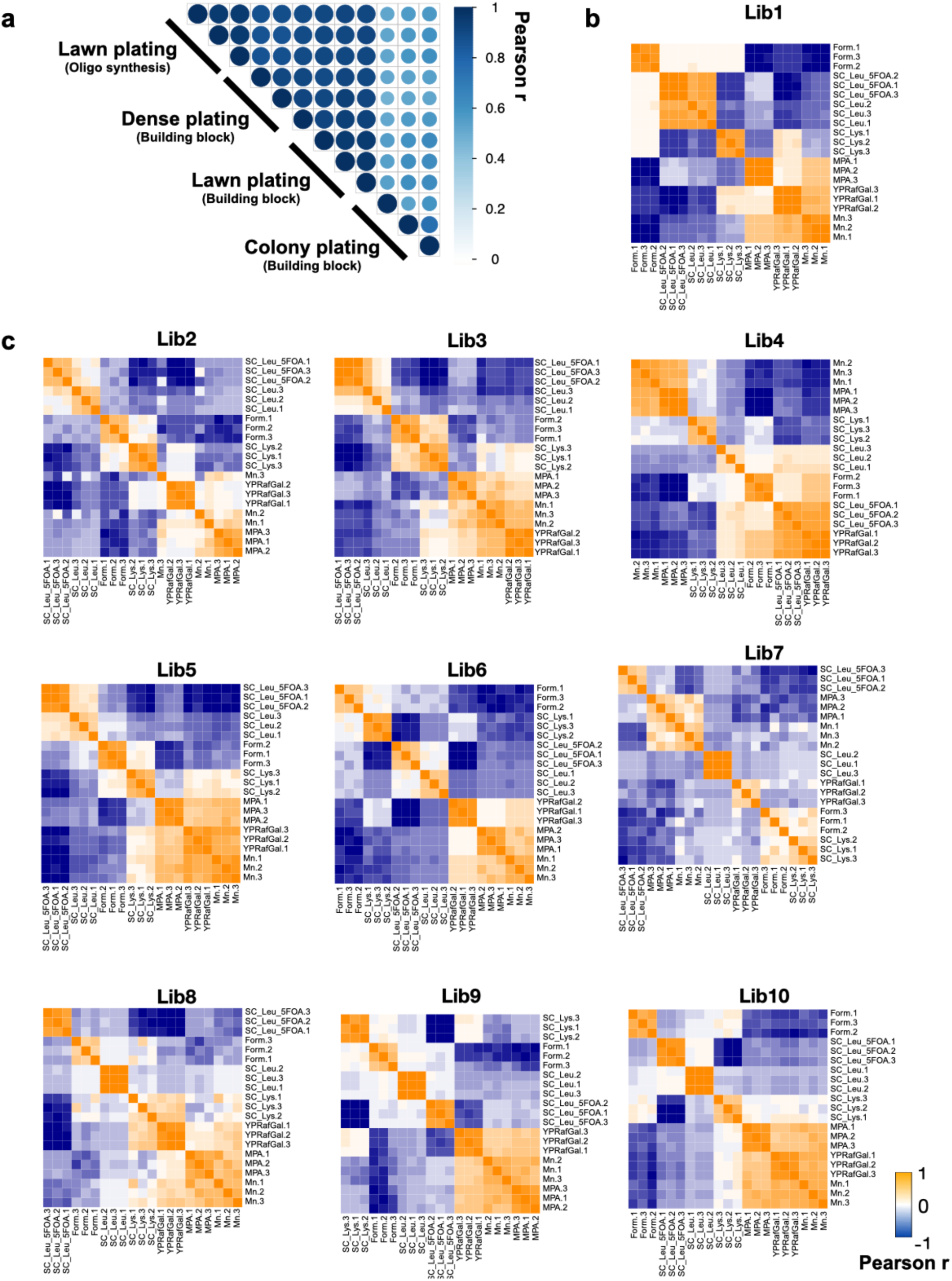
Pol II deep mutational scanning is highly reproducible. **a.** Single mutant growth fitness from mutants in libraries constructed from synthesized oligos correlated well with our previous library constructed by a random building block approach when plating conditions were the same. Qiu *et al*^17^ plated at a lower density (colony plating) that we speculated added noise to the analysis. When plating densely (“dense” and “lawn” conditions) our new and old libraries showed highly reproducible fitness determinations for single mutants. **b-c.** Biological replicates for each library showed high reproducibility for all conditions. Pearson correlation of each library was calculated with three replicates for viable mutant fitness on all selective conditions.

**Extended Data Fig.2.**
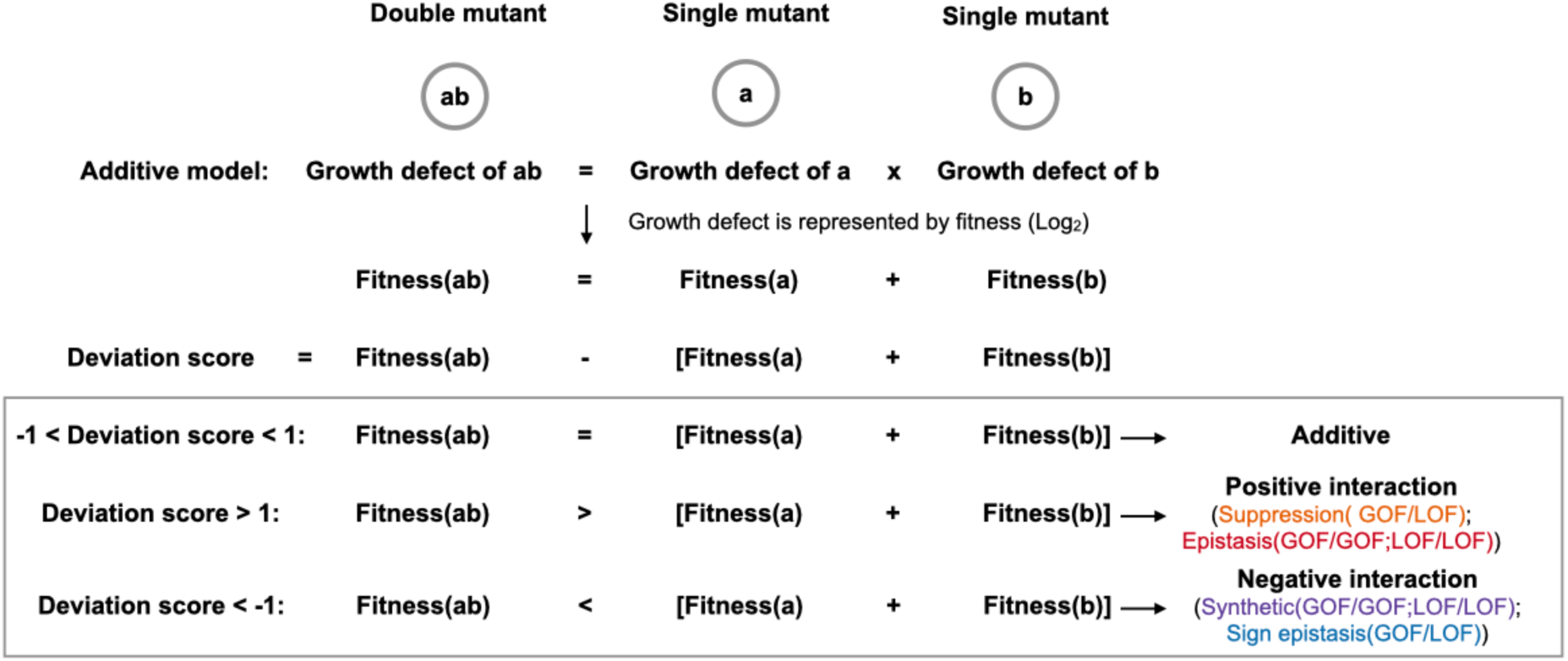
Detection of functional interactions by deviation score. For a pseudo double mutant *ab*, the difference between its observed fitness (*ab*) and expected fitness (*ab*) adding the fitness of two constituent single mutants (*a* and *b*) determines the type of interaction between the two mutants. Positive or negative interactions are determined if the deviation score is greater than 1 or smaller than –1. Specific epistatic interactions are further distinguished from general suppression or synthetic sick or lethal interactions using predicted mutant catalytic defect classes (GOF or LOF).

**Extended Data Fig.3.**
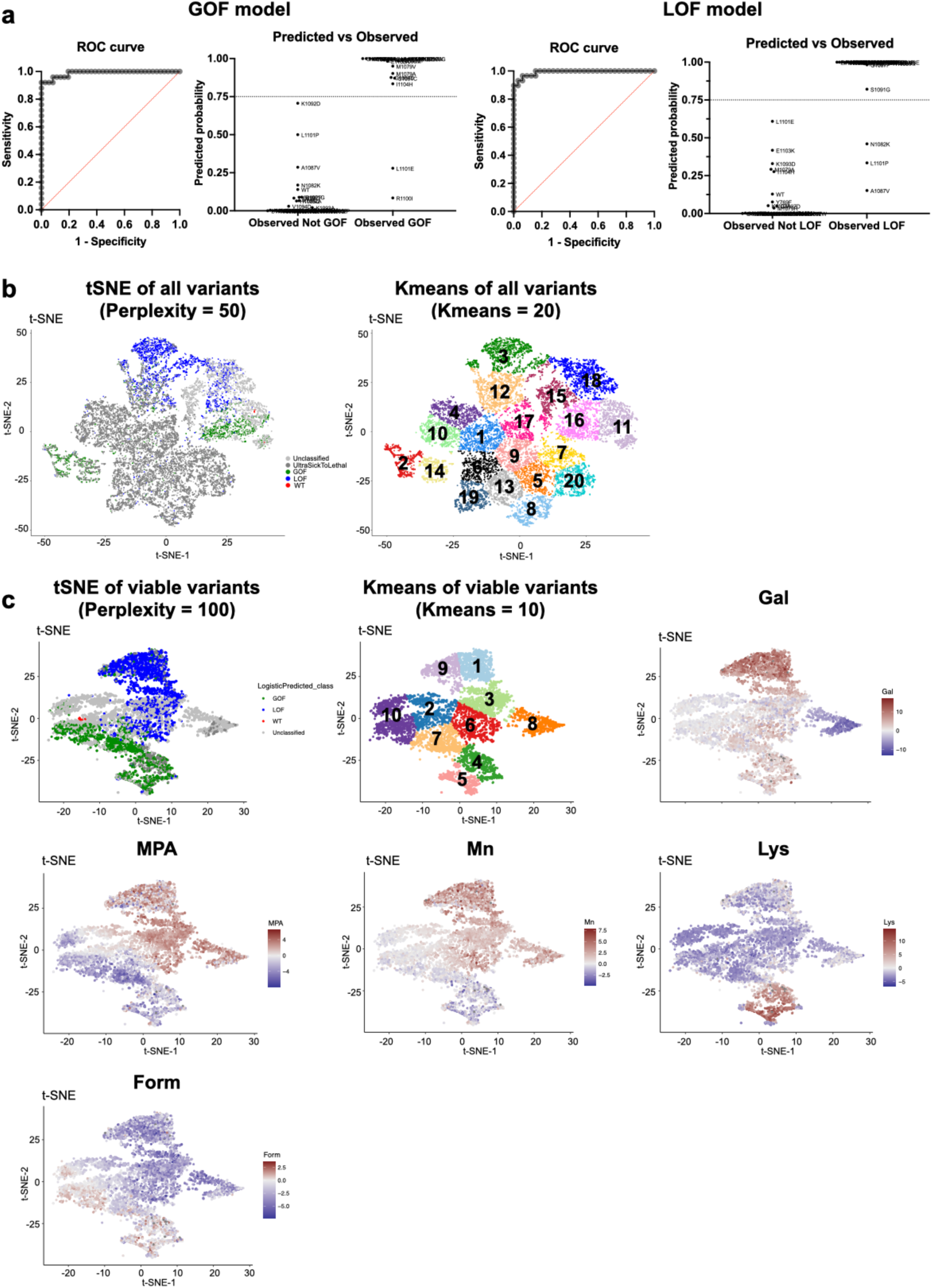
Classification of mutant catalytic defects with machine learning algorithms. **a.** ROC curves for two multiple logistic regression models used to determine mutant catalytic class. Using 65 mutants with validated in vitro determined catalytic defects and conditional growth fitness measured in our experiment, we trained two models to classify variants as GOF or LOF. The GOF AUROC is 0.9889 (P ≤ 0.0001), whereas the LOF ROC is 0.9914 (P ≤ 0.0001). The predicted vs. observed graphs display the predicted probability of 65 known mutants would be GOF or LOF. The threshold we used to determine GOF or LOF mutations is shown by lines at 0.75. Details of the models are in Supplemental Table 6. Among the 6054 viable mutants, 1390 were classified as GOF (22.96%), 1702 as LOF (28.11%), and 2962 remained unclassified (48.93%). **b.** Left: t-SNE projection of all mutants (n=15174) with perplexity = 50. Right: k-means cluster of all mutants with 20 clusters. The t-SNE and k-means projections suggest GOF are in 3 clusters (cluster 2, 14, and 16), LOF are in 2 clusters (cluster 3 and 18), and unclassified mutants are in 2 clusters (11 and 15). Most ultra-sick/lethal mutants (fitness <= -6.5) are projected together into 13 clusters, likely due to significant noise from low read counts across conditions. **c.** Feature plot of viable mutations in t-SNE and k-means projections (n=6054). Ultra-sick/lethal mutations were removed and the viable mutants were projected with t-SNE (perplexity = 100) and K-means (10 clusters). GOF were grouped into 4 clusters (4, 5, 7 and 10) and LOF were in 4 clusters (1, 3, 6, and 9). Each spot in the projection represents a mutant and it is colored based on the fitness of the mutant in selective conditions. GOF and LOF mutants in different clusters are related to various phenotype patterns. GOF clusters 7 and 10 are defined by strong MPA^S^, while clusters 4 and 5 show slight MPA^S^, Gal^R^, Mn^S^, but strong Lys^+^. Slight Form^S^ is a common feature across four GOF clusters. LOF clusters 3 and 6 show slight Mn^R^, while clusters 1 and 9 are strongly Mn^R^ and Gal^R^. There are three common features in all LOF clusters: MPA^R^, Form^S^, and Lys^-^. Cluster 8, which mostly contain unclassified mutants, appear defined by Gal super sensitivity, indicating a potential specific defect defining this cluster.

**Extended Data Fig.4.**
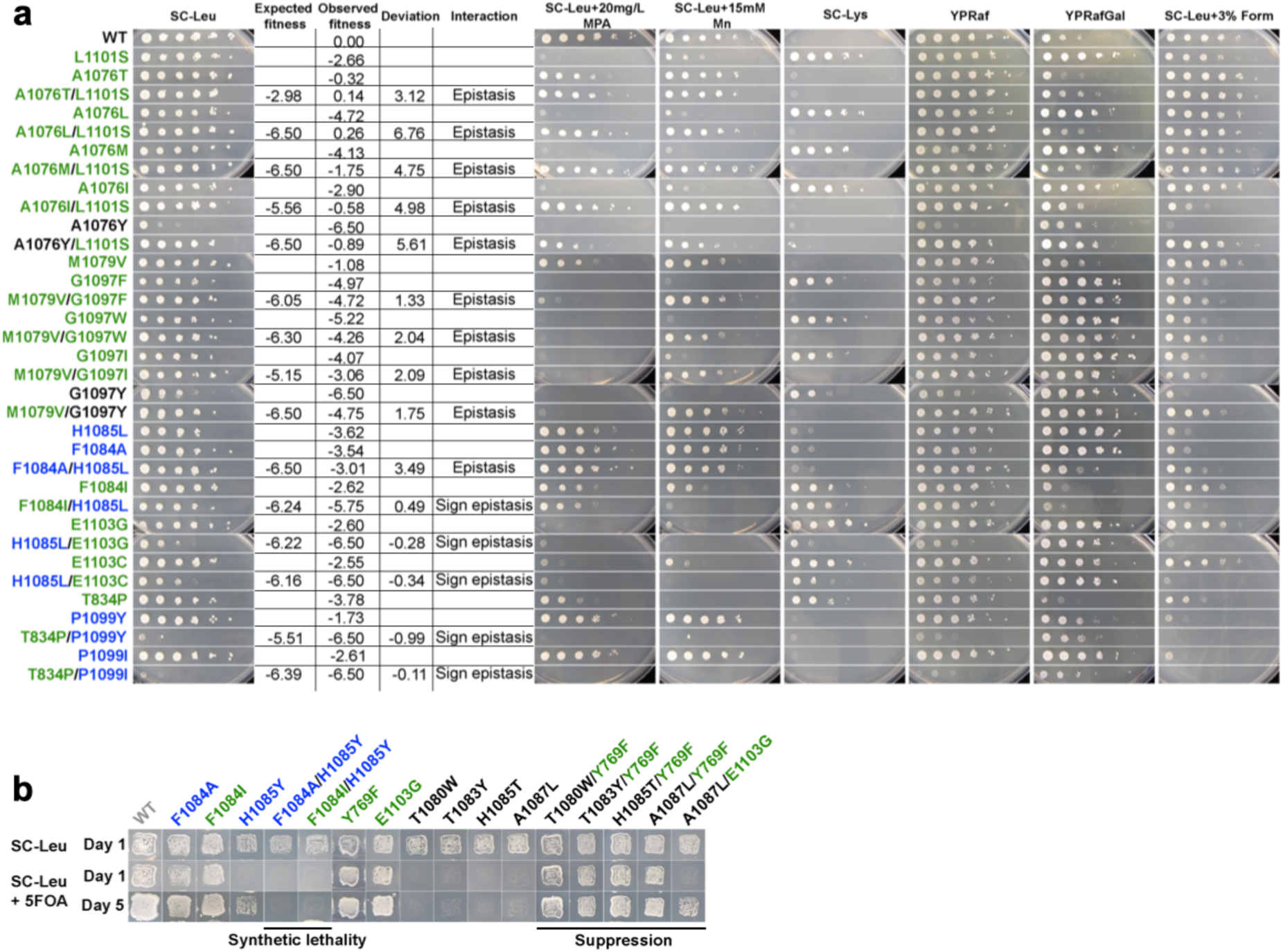
Genetic interactions detected with spot assay and patch assay are consistent with high-throughput results. To verify the genetic interactions observed in high throughput experiment, we picked 50 mutants to measure the genetic interactions with spot assay (**a**) or patch assay (**b**). They are consistent. The fitness, deviation scores and corresponding genetic interactions measured in high throughput experiment are indicated in **a.** Specifically, the labeled observed fitness indicates observed general growth fitness that was measured on SC-Leu+5FOA.The names of GOF mutants are labeled with green color, LOF mutants are labeled with blue, ultra-sick to lethal mutants are labeled with black color.

**Extended Data Fig.5.**
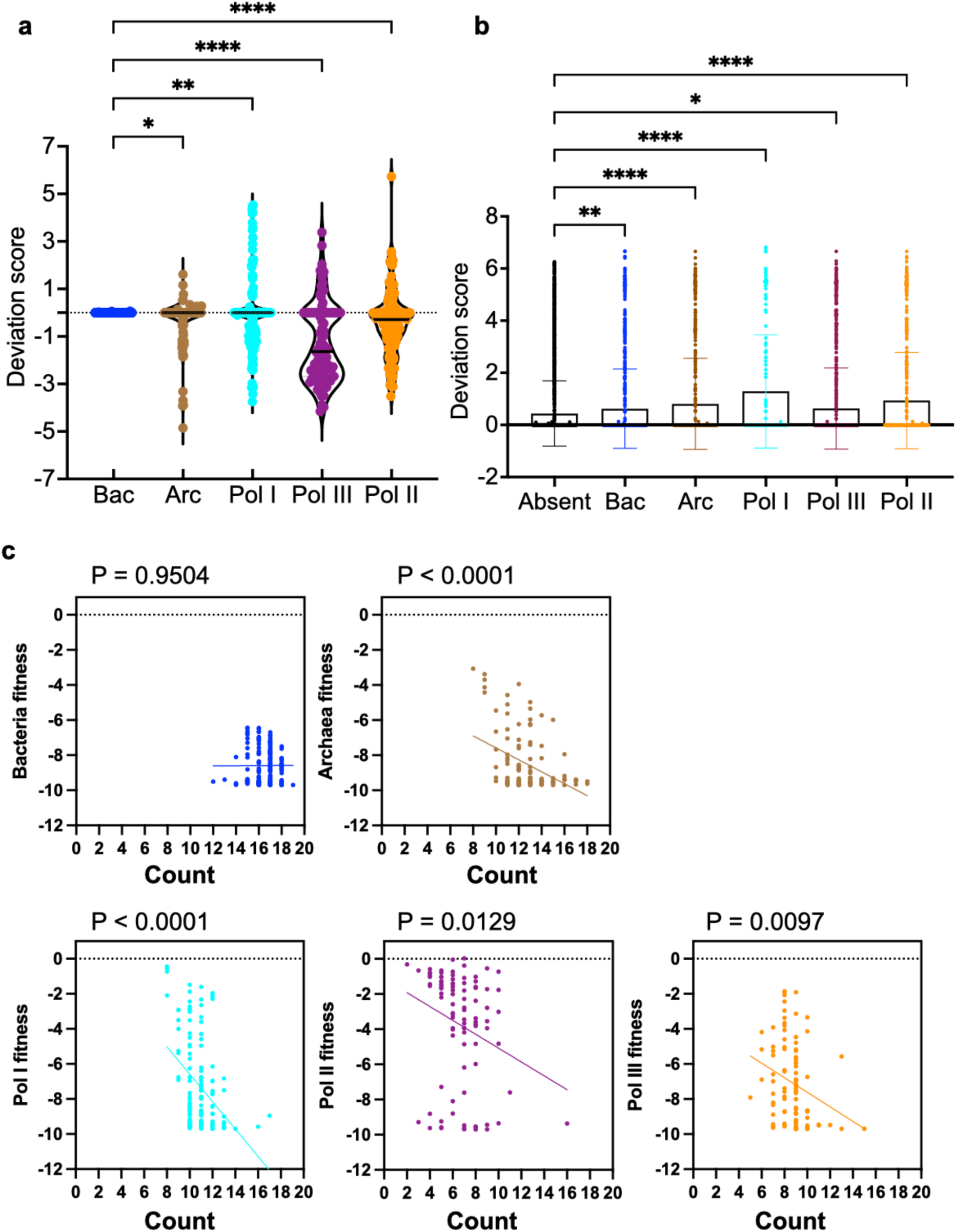
Contextual epistasis affects fitness of TL haplotypes. **a.** Distributions of deviation scores of the TL haplotypes in each group. **b.** Comparison of the mean deviation scores of lethal single substitutions that are present in different species and those that are absent in any species. Standard deviation values are also shown in the bar plot. ANOVA multiple comparison was applied to compare the mean deviation score of the “Absent” group to each of the other groups and significant levels (P < 0.05) are shown in the figure. **c.** An xy-plot of evolutionary observed TL haplotypes fitness versus the numbers of substitutions in the haplotypes. Simple linear regression was performed for each plot. Bacteria fitness vs count: Y = 0.004267*X – 8.660, r^2^=2.152e-005. Archaea fitness vs count: Y = -0.3406*X - 4.175, r^2^=0.1568. Pol I fitness vs count: Y = -0.7818*X + 1.235, r^2^=0.1521. Pol II fitness vs count: Y = -0.3943*X – 1.132, r^2^=0.06535. Pol III fitness vs count: Y = -0.4148*X – 3.468, r^2^=0.06984. The P values of the slopes are labeled.

**Extended Data Fig.6.**
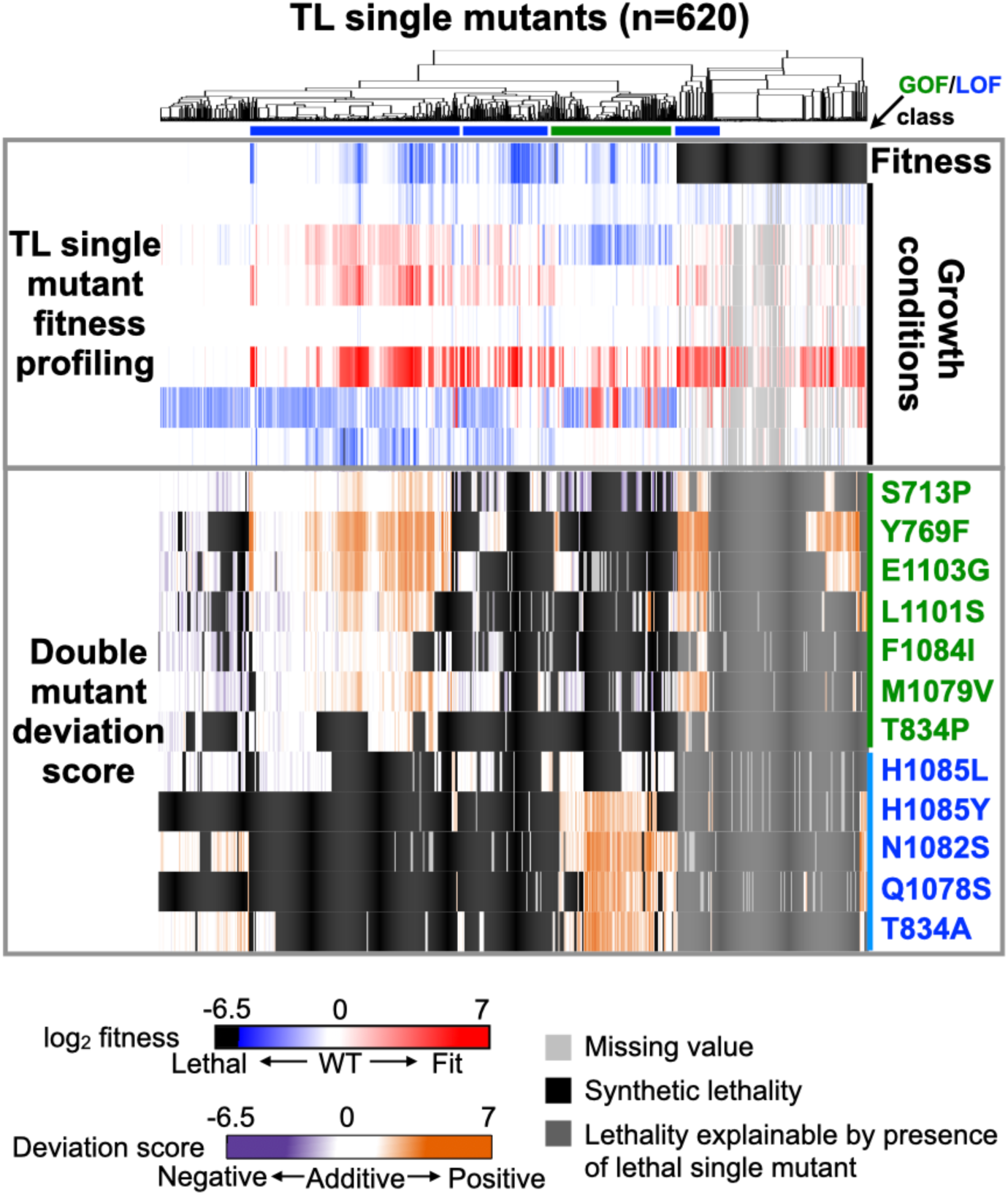
The functional interaction landscape of probe mutants. The functional interaction landscape is shown as a heatmap. The upper part of the heatmap shows all Pol II TL single mutant growth fitness profiling across several phenotypes and the single mutants were ordered by hierarchical clustering with Euclidean distance. The lower part of the heatmap shows double mutant deviation scores where a colored block at the interaction of x and y coordinates indicates deviation score of the double mutant.

**Extended Data Fig.7.**
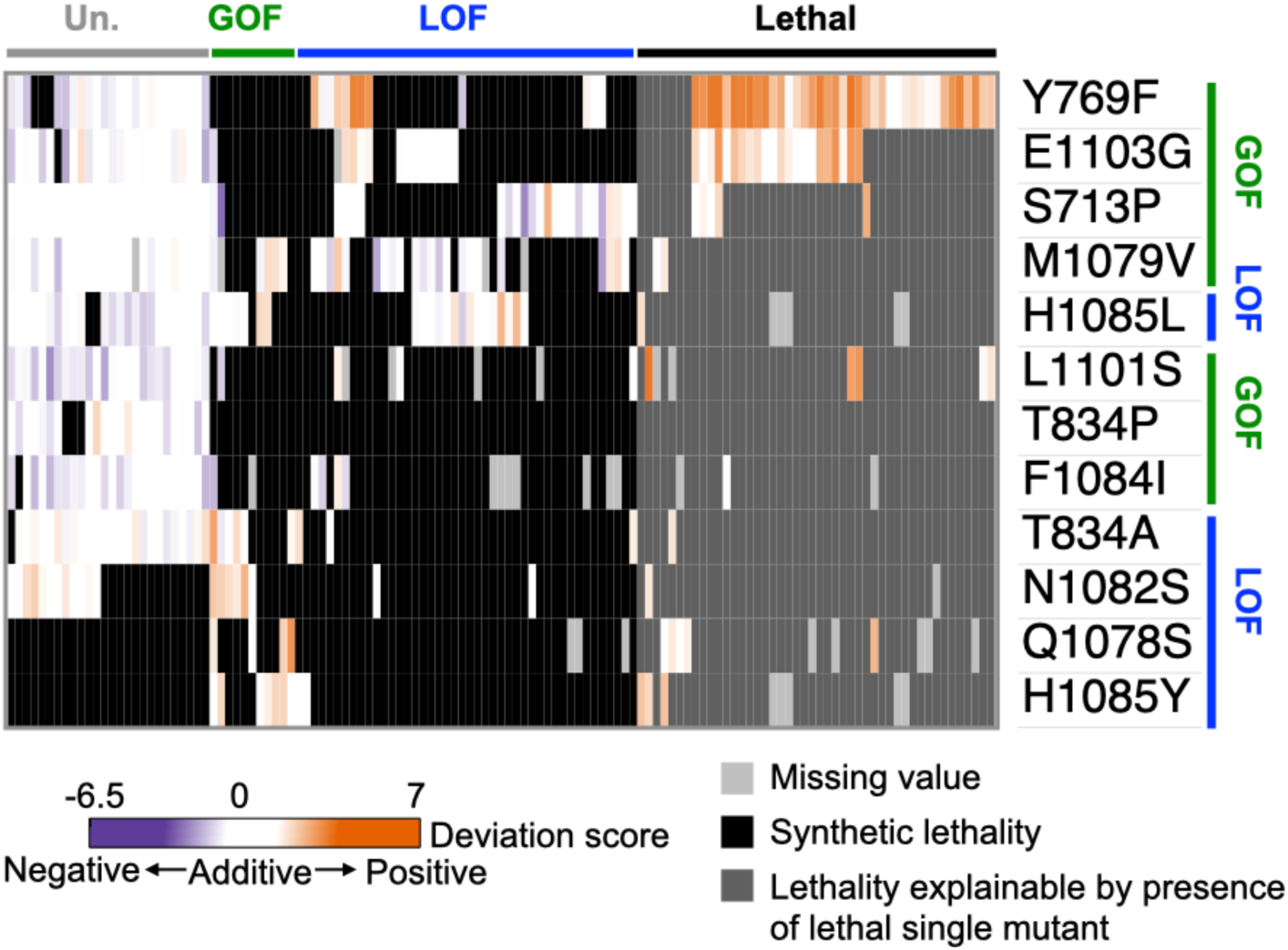
Allele-specific interactions. Unique interactions observed between TL substitutions and probe mutants. For each substitution, we analyzed the interquartile range (IQR) of their deviation scores with all probe mutants. Any substitution with deviation score(s) outside of the IQR were extracted and called as unique interaction(s). 127 substitutions with unique interactions with probe mutants were found out of 620 and are shown in the heatmap. The heatmap was hierarchical clustered with Euclidean distance for both rows and columns.

**Extended Data Fig.8.**
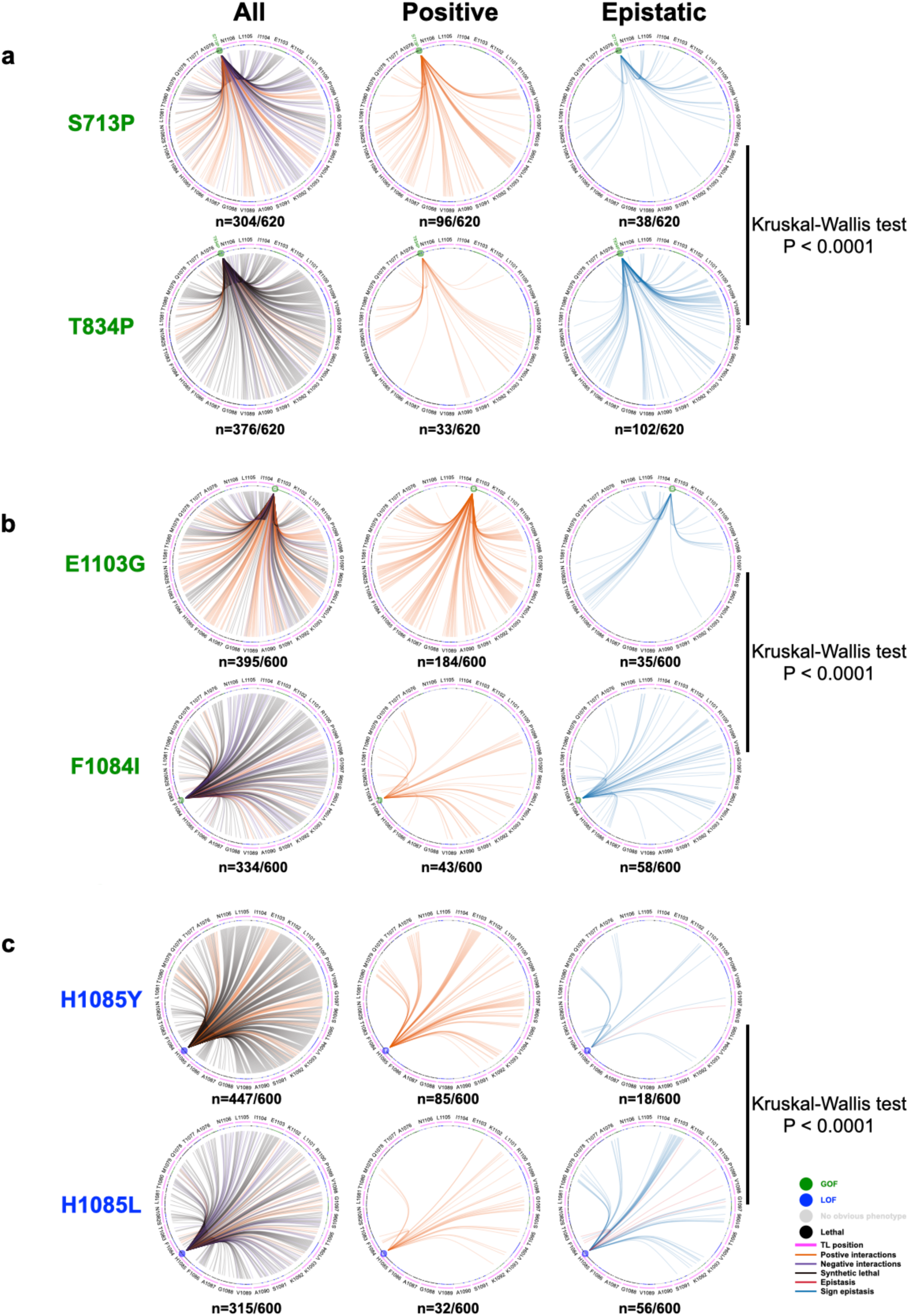
Interaction networks of selected probe mutants. The TL is shown in circle with WT residues and positions labeled. All 20 substitutions of each TL residue are represented by a magenta arc under each WT residue, with tick marks representing individual substitutions at that position and are colored by mutant class. Comparison of interaction networks between S713P and T834P (**a**), E1103G and F1084I (**b**) and H1085Y and H1085L (**c**) showed the differences are significant (P < 0.0001). The comparisons were performed with Kruskal-Wallis test with P-value correction with Dunn’s multiple comparisons test (Supplemental Table 7).

**Extended Data Fig.9.**
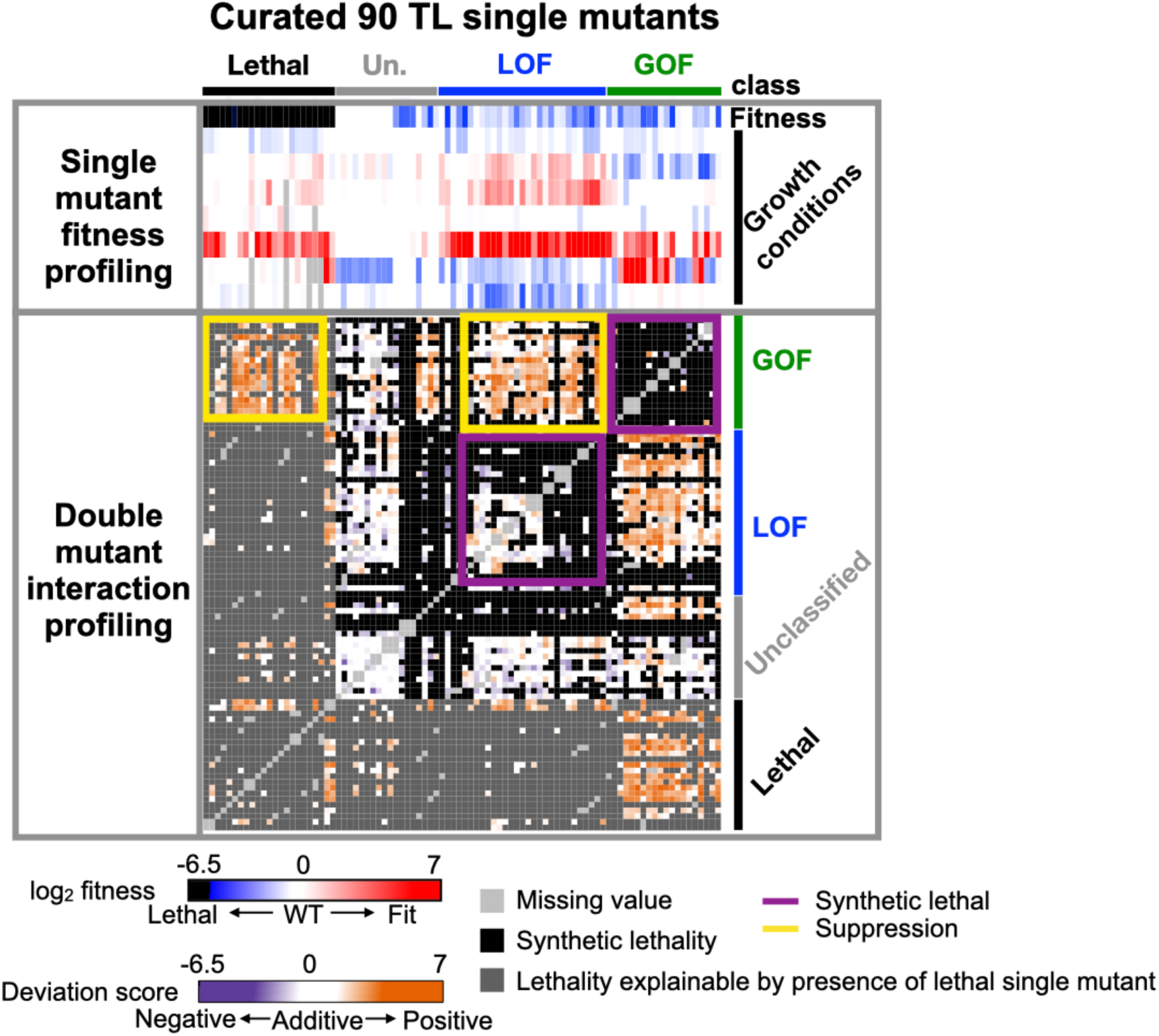
The intra-TL functional interaction landscape. The intra-TL functional interaction landscape is shown as a heatmap. Annotations at the top and right indicate the 90 curated single mutants and their predicted phenotypic classes from multiple logistic regression modeling. The upper part of the heatmap shows single mutant growth fitness profiling across multiple phenotypes ordered by groups predicted with logistic regression models. The lower part of the heatmap shows double mutant deviation scores where a colored block at the interaction of x and y coordinates indicates deviation score of the double mutant.

**Extended Data Fig.10.**
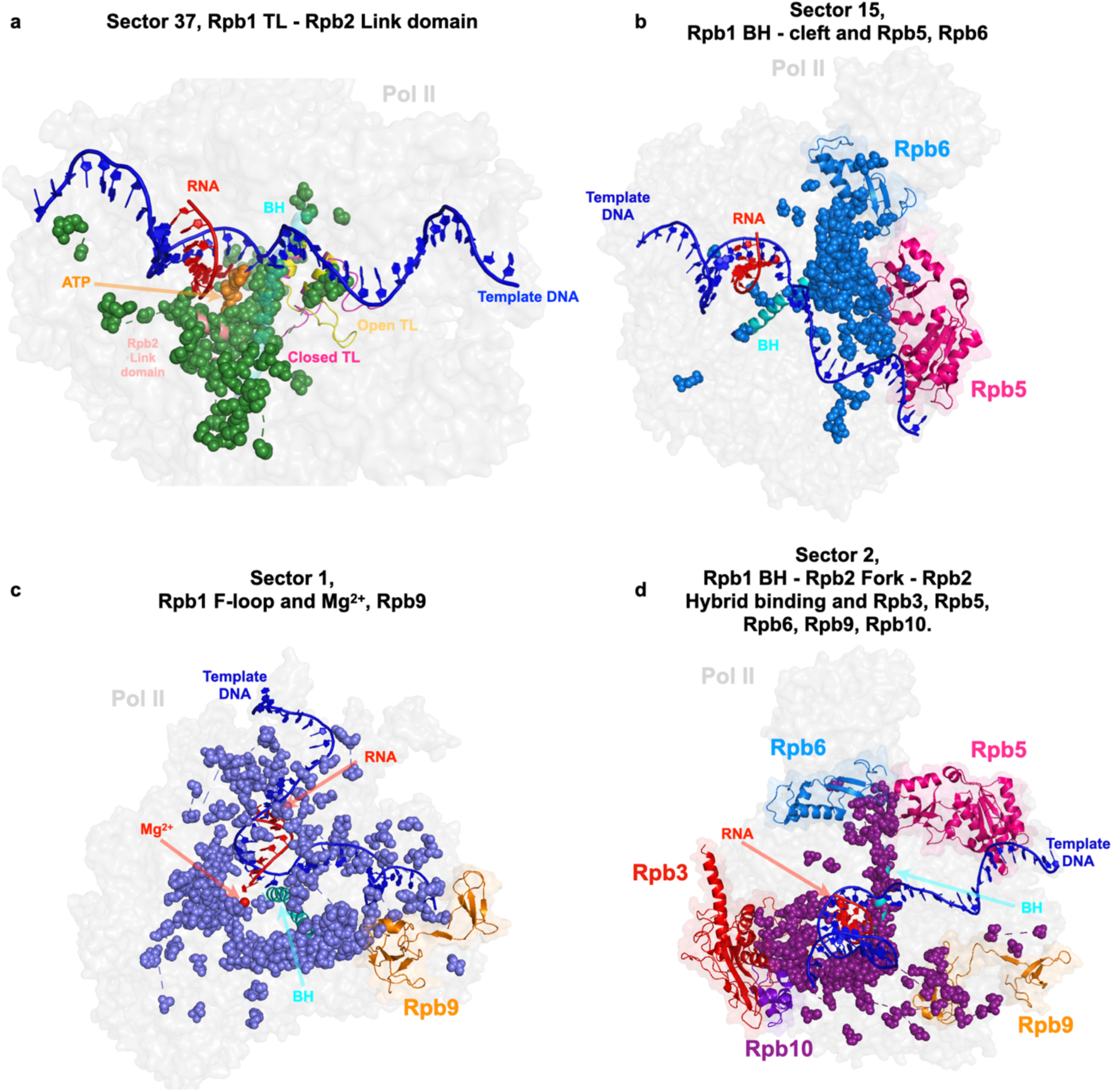
Coevolutionary sectors of Pol II active site residues extend to the surface of Pol II, forming connections with other Pol II subunits. **a**. The coevolutionary sector of Rpb1 TL and Rpb2 link domain residues surrounds the substrate ATP, indicating its potential role in catalysis. The sector is mapped on the yeast Pol II structure with TL in open state (PDB: 5C4X) and closed state (PDB: 8U9R). **b**. Residues in sector 15 connect with Rpb5 and Rpb6. **c.** Residues on sector 1 connect with Mg^2+^ and Rpb9. **d.** Residues in sector 2 connect with Rpb5, Rpb6, Rpb3, Rpb9 and Rpb10. These sectors are mapped on the yeast Pol II structure (PDB: 5C4X)^42^.

