## Supplemental Figures for "Widespread epistasis shapes RNA Polymerase II active site function and evolution"

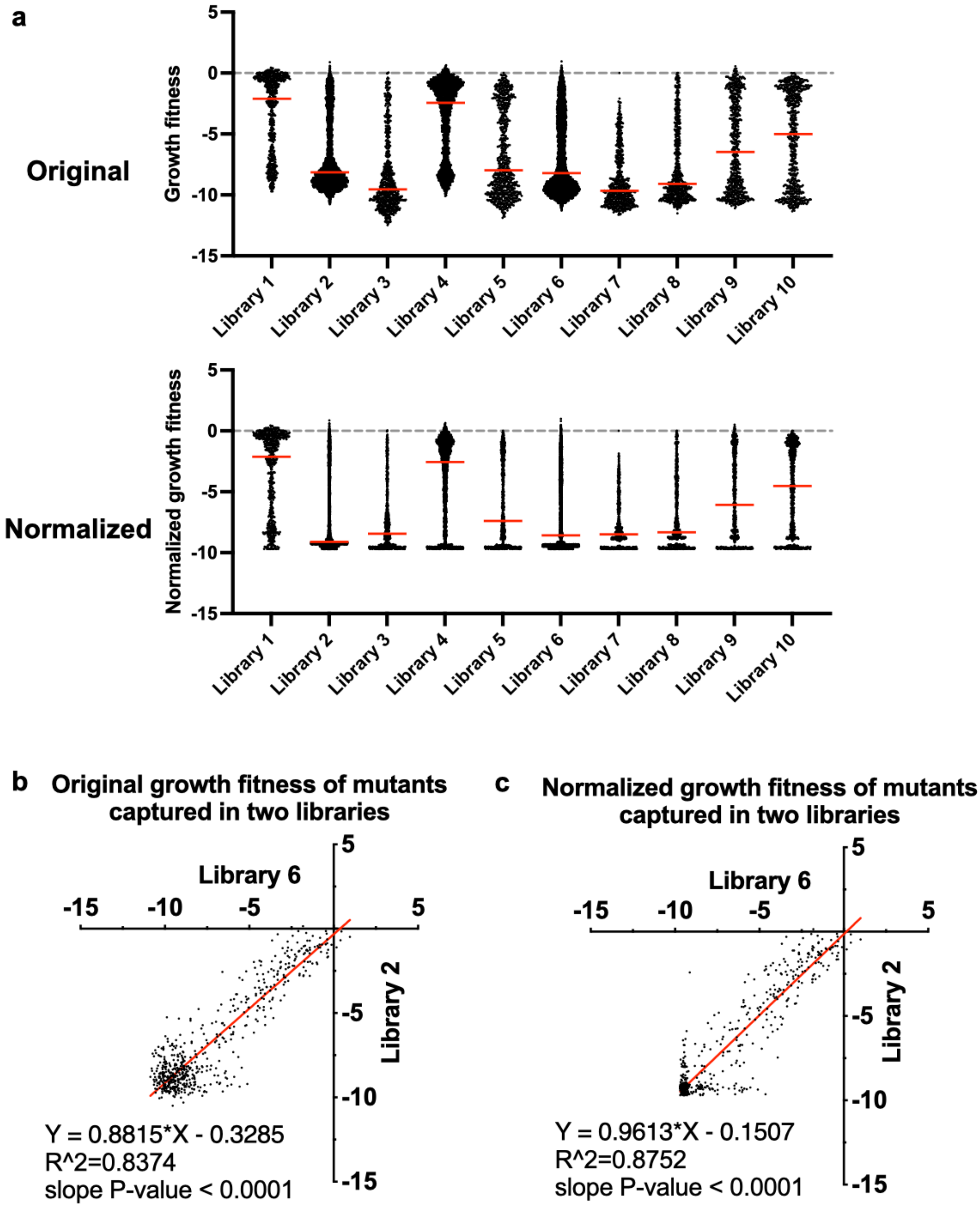


**Figure S1. Min-Max normalization uniformed the fitness level of lethal mutants from different libraries without disturbing the median of mutant fitness. a.** Library growth fitness distributions before and after normalization. Upper panel: The growth fitness distributions of libraries. The lowest fitness levels (fitness of lethal mutants) are different among libraries. To uniform various lowest fitness levels, we applied Min-Max normalization to minimize library effects on fitness ranges (See **Methods** for details). Lower panel: Libraries fitness distributions after normalization. The lethal mutant fitness levels of libraries were normalized to the same level while the median fitness for each library was not affected by the normalization. **b-c.** XY-plots showing the original fitness of mutants captured in two different libraries (n=586) (**b**). These mutants present in two libraries showed improved correlation between measurements upon normalization (**c**).

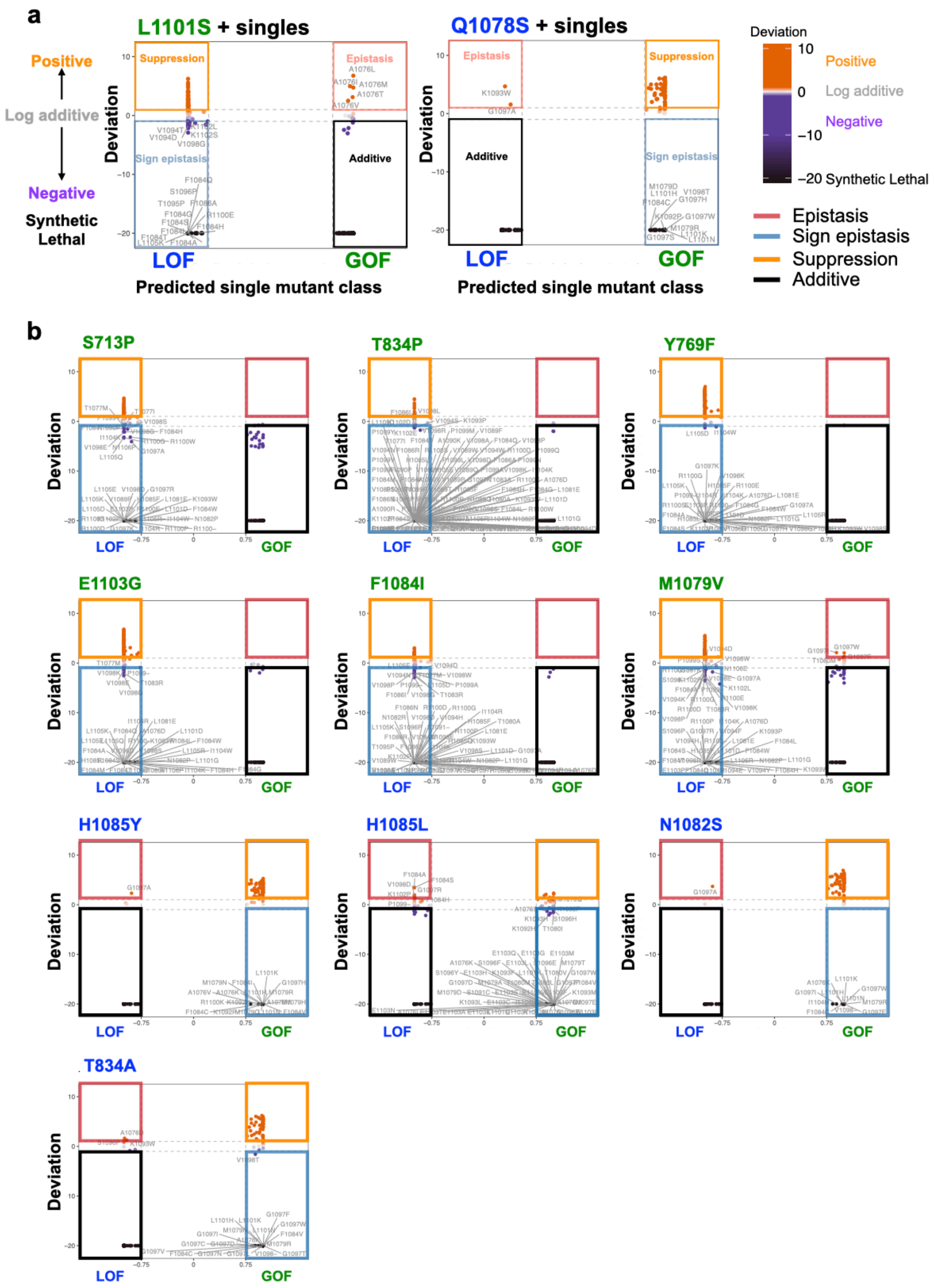

a
L1101S + singles
Q1078S + singles
Positive
Log additive
Negative
Synthetic Lethal
Suppression
Epistasis
Sign epistasis
Additive
Deviation
LOF
GOF
Predicted single mutant class
Deviation
10
0
−10
−20 Synthetic Lethal
Positive
Log additive
Negative
Epistasis
Sign epistasis
Suppression
Additive
b
S713P
T834P
Y769F
E1103G
F1084I
M1079V
H1085Y
H1085L
N1082S
T834A

**Figure S2. Identifying TL substitutions that interact with the probe mutants. a.** Identification of epistasis and suppression within positive interactions, and sign epistasis and synthetic sickness/lethality within negative interactions in two probe mutants, L1101S and N1082S. The deviation score of combinations (y-axis) between probe mutants and TL GOF or LOF single mutants were plotted versus the predicted probability of single mutants being GOF or LOF (x-axis). **b.** The scatter plots for distinguishing interactions of the other 10 TL probe mutants.

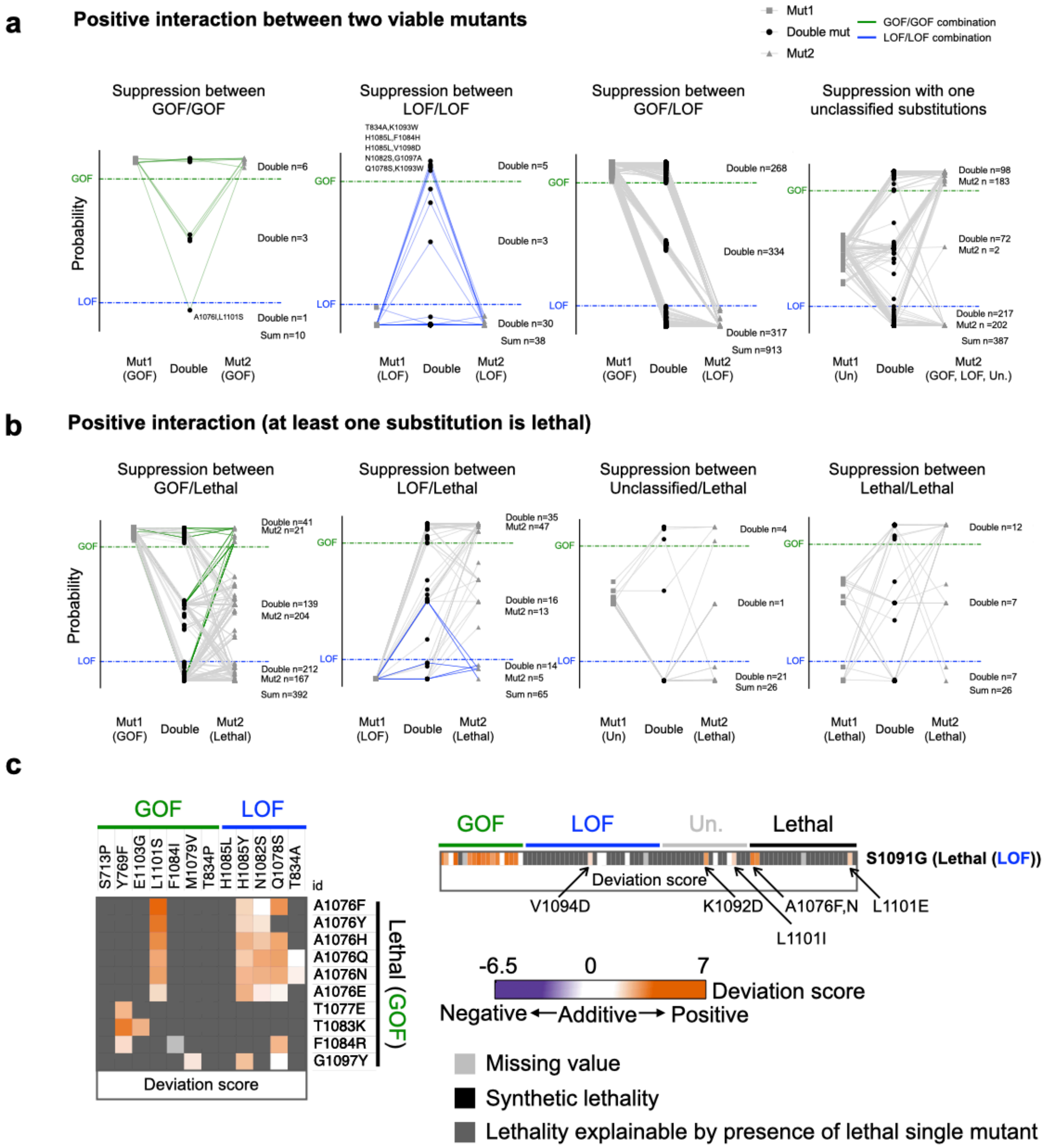


### Ratio of observed strong or weak epistasis

| | Strong epistasis (\|d\| > 2) | Weak epistasis (1< \|d\| < 2) |
|---|---|---|
| Pairwise Intra-TL doubles | 0.15 | 0.06 |
| Target Intra-TL doubles | 0.15 | 0.09 |
| Target Pol II-TL doubles | 0.17 | 0.10 |
| Reported | 0.05 | 0.3 |

**Figure S3. Discrimination of regular epistasis from sign epistasis. a.** The phenotypic classes of double mutants consist of two viable single substitutions with positive interactions. Four plots show four kinds of combinations respectively. For each plot, the predicted GOF or LOF probabilities of a double mutant and two constituent single mutants are shown in Y-axis. The double mutant and two constituent single mutants are shown in X-axis in the order of the first constituent mutant (Mut1), the double mutant (Double), the second constituent mutant (Mut2). The double mutant and two constituent mutants are connected with lines for each pair of combinations. The numbers of double mutants belonging to GOF (top), unclassified (middle), or LOF (bottom) are labeled. GOF and LOF probability threshold are labeled with dashed lines. **b.** The phenotypic classes of double mutants consist of one viable and one lethal single substitution, or two lethal single substitutions with positive interactions. The arrangements of plots are similar to **a**. **c.** The heatmaps of lethal GOF substitutions suppressed by GOF targets (left) and lethal LOF substitutions suppressed by LOF targets (right). **d.** The fraction of strong and weak interactions we observed in double mutants compared with the ratio reported in other studies [57,86-89].

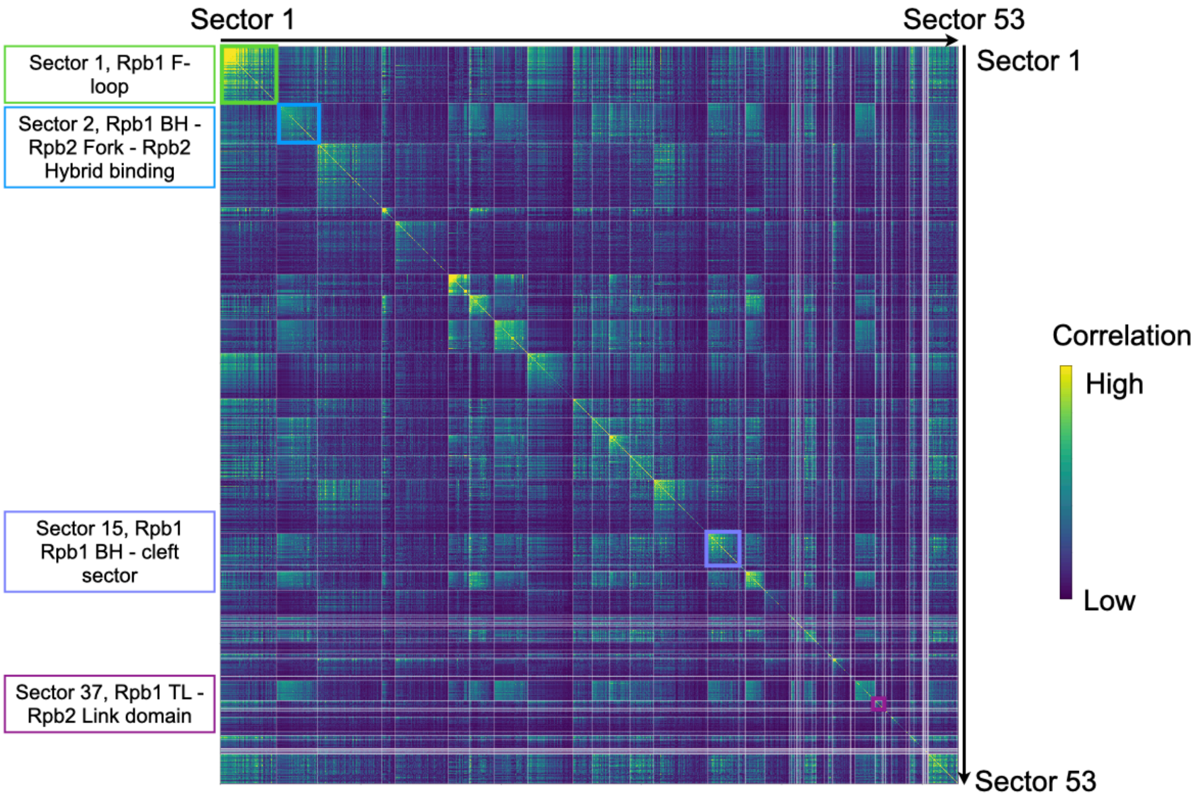


**Figure S4**. **Rpb1/Rpb2 coevolutionary residue sectors identified by Statistical Coupling Analysis (SCA).** 53 significant and independent sectors are shown in a heatmap with correlation score calculated from the statistical coupling analysis. Four major sectors containing Pol II active site residues are labeled on the left of the heatmap. Statistical coupling analysis was applied to a Multiple Sequence Alignment (MSA) of 283 Rpb1/Rpb2 homologs.

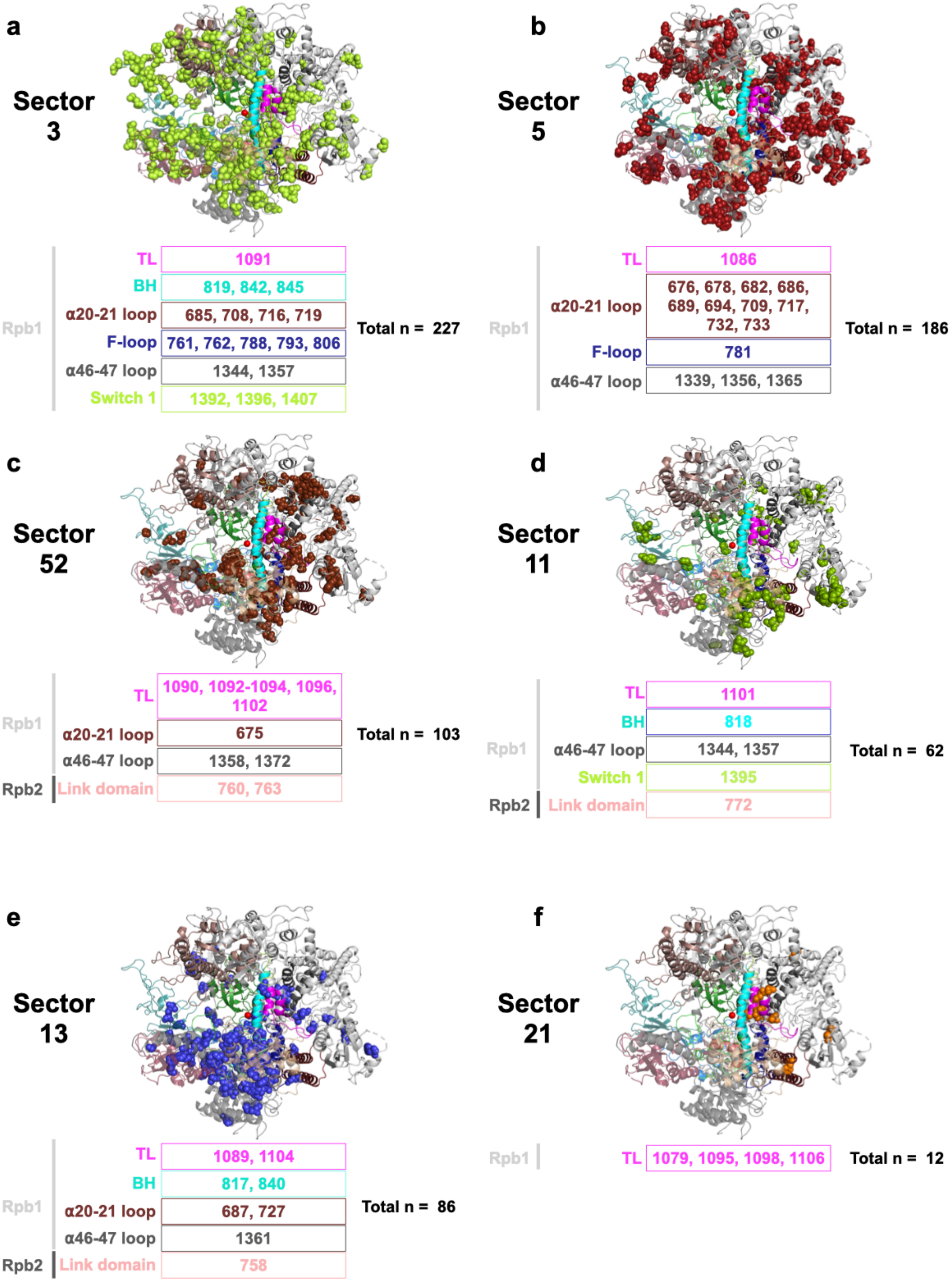

a
Sector 3
Rpb1
TL
1091
BH
819, 842, 845
α20-21 loop
685, 708, 716, 719
F-loop
761, 762, 788, 793, 806
α46-47 loop
1344, 1357
Switch 1
1392, 1396, 1407
Total n = 227
b
Sector 5
Rpb1
TL
1086
α20-21 loop
676, 678, 682, 686, 689, 694, 709, 717, 732, 733
F-loop
781
α46-47 loop
1339, 1356, 1365
Total n = 186
c
Sector 52
Rpb1
TL
1090, 1092-1094, 1096, 1102
α20-21 loop
675
α46-47 loop
1358, 1372
Rpb2
Link domain
760, 763
Total n = 103
d
Sector 11
Rpb1
TL
1101
BH
818
α46-47 loop
1344, 1357
Switch 1
1395
Rpb2
Link domain
772
Total n = 62
e
Sector 13
Rpb1
TL
1089, 1104
BH
817, 840
α20-21 loop
687, 727
α46-47 loop
1361
Rpb2
Link domain
758
Total n = 86
f
Sector 21
Rpb1
TL
1079, 1095, 1098, 1106
Total n = 12

**Figure S5**. **Other coevolutionary networks involving Pol II active site residues.** In addition to the four major sectors, Pol II active site residues, along with residues in Rpb1/Rpb2, form additional coevolutionary sectors (**a-f**). Residues within the Pol II active site domains are labeled. These sectors are mapped on the yeast Pol II Rpb1/Rpb2 structure (PDB: 5C4X)[42].

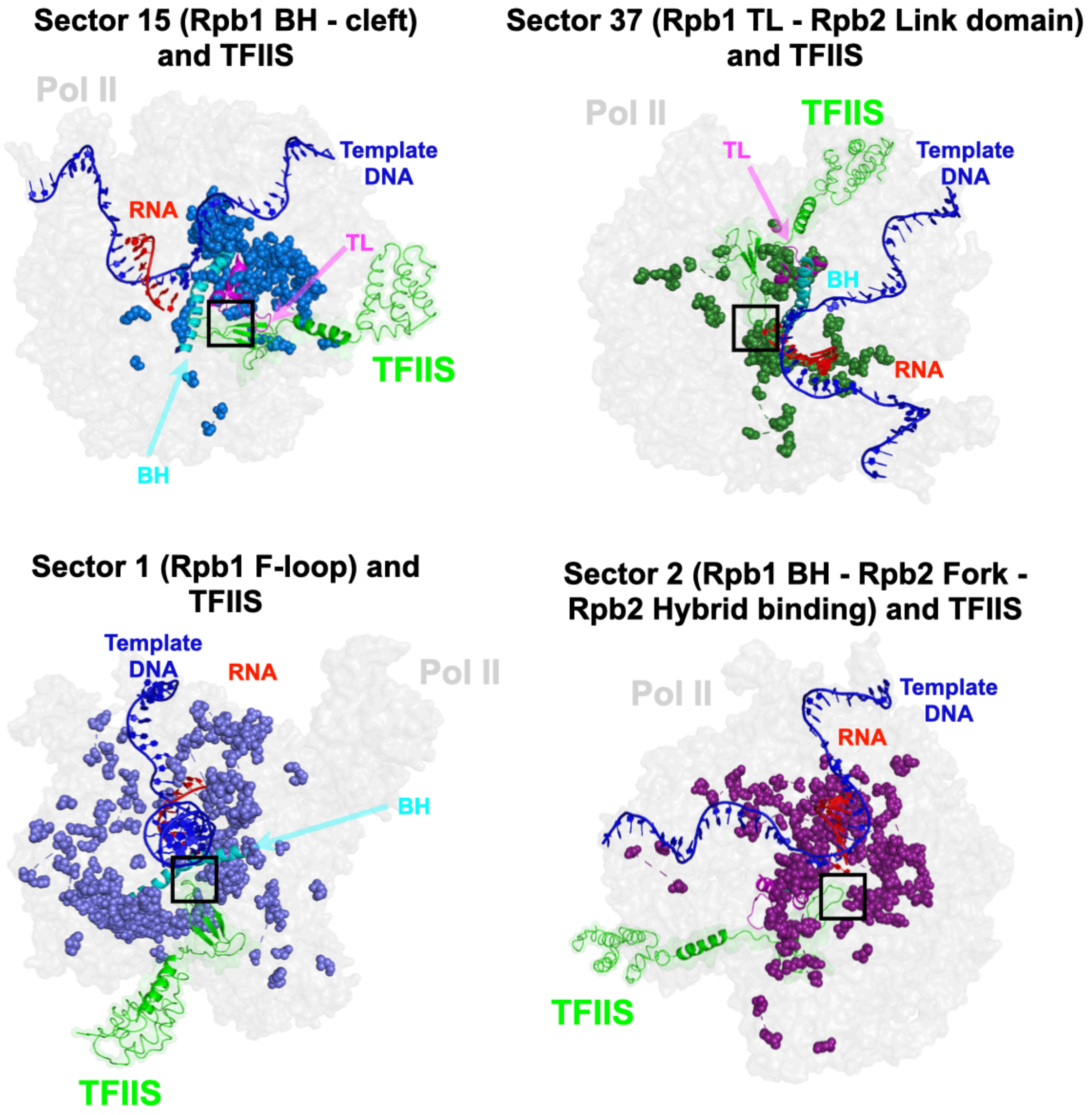


**Figure S6. Coevolutionary residues of four major sectors in Pol II active site interact with TFIIS.** Coevolutionary residues are mapped on the yeast Pol II structure (PDB:5C4X) and aligned with TFIIS structure (PDB:1Y1V). The interacting residues are highlighted with black box respectively.

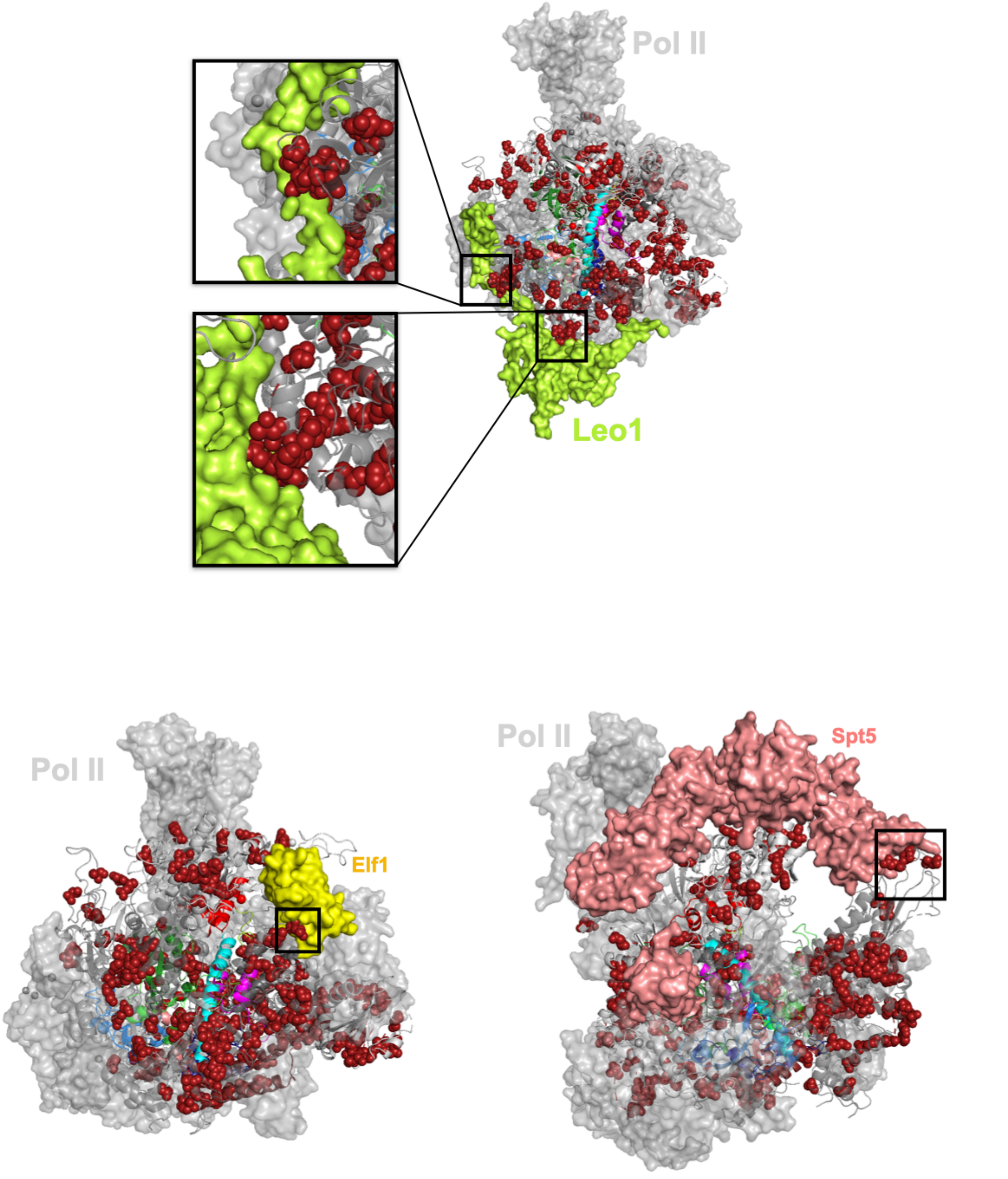


**Figure S7. Residues in Pol II active site coevolutionary sectors extend to the surface of the enzyme and interact with transcription factors.** Coevolutionary residues in sector 5 (**Fig. S16b** in the manuscript) are mapped on the yeast Pol II structure (PDB:5C4X) and aligned with the structures of transcription factors Leo1, Elf1, and Spt5 (PDB:7xn7). The interacting residues are highlighted with black box respectively.

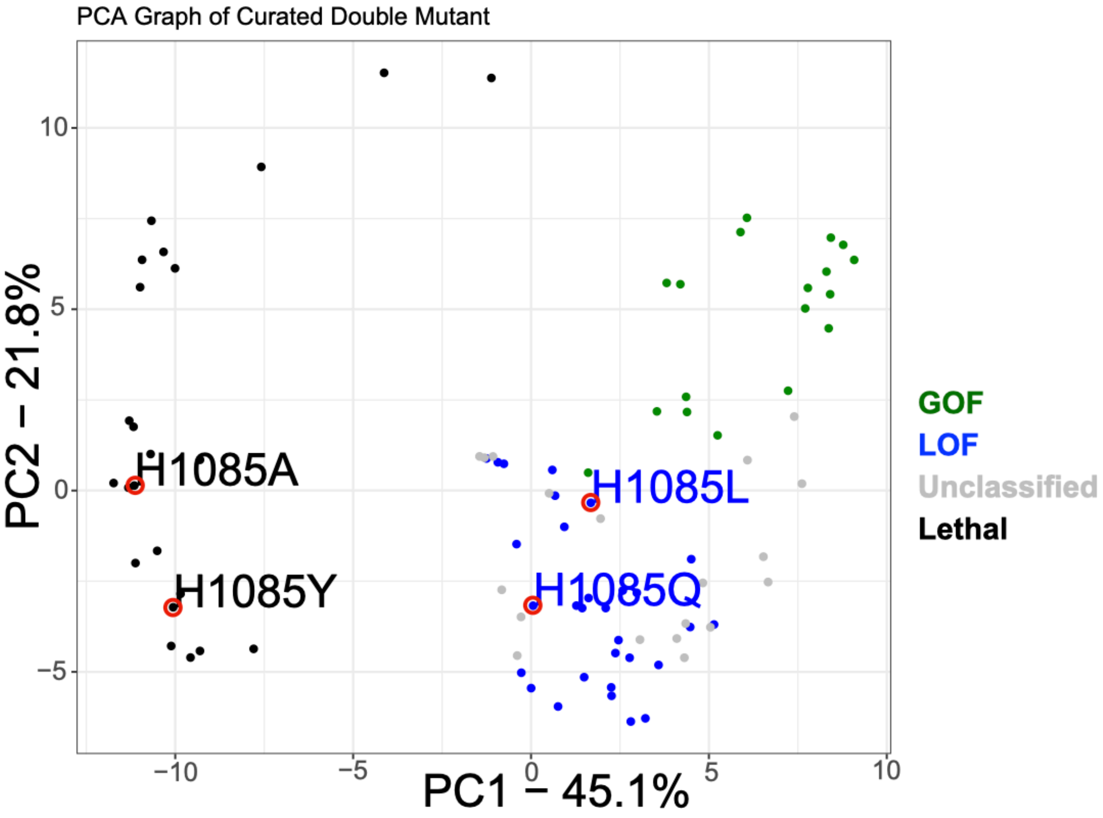


**Figure S8**. **Four H1085 substitutions are different in some ways. A.** Principal component analysis (PCA) with double mutant deviation scores of all curated TL single mutant substitutions, which are represented with colored dots. GOF mutants are in green, LOF mutants are in blue, unclassified mutants are in grey and lethal mutants are in black. Four H1085 substitutions are labeled and assigned with a red circle to make them visible in the plot.
