## Supplemental Methods for "Widespread epistasis shapes RNA Polymerase II active site function and evolution"

**Supplemental method 1: High efficiency large scale chemical yeast transformation protocol**

We optimized the Gietz protocol[1], which yields 5-15 x $10^5$ colonies with 1μg of DNA and 10 × $10^8$ cells. However, transformation efficiency may vary with different DNAs and yeast strains.

Day 1:
Set up 5mL cultures from a single colony in YPD.

Day 2:

1. Dilute ~400-500μL of saturated culture in 50mL of YPD (210rpm, 30 °C). The cells were grown for ~3-4 hours until the concentration of the culture reached 2 × $10^7$ cells/mL (count the cells).

2. The cells were washed in 25 ml of sterile water and resuspended in 1 ml sterile water (volume needs to be adjusted to make the final concentration 1 × $10^9$ cells/mL).

3. Boil ssDNA for 10mins before use, and the boiled ssDNA was kept on ice for the entire duration.

4. Aliquot of 100μL ($10^8$ cells) were centrifuged at top speed for 30s, discarded the supernatant was discarded, and the cells were resuspended in the transformation mix (240μL 50% PEG 3500, 36μL 1M LiOAc, 10μL ssDNA, 74μL DNA with water to make 100ng in total, 360μL in total).

5. The mixture was then incubated at 30 °C for 30 min.

6. Tape the tubes to a 30 °C wheel for 30 min of incubation.

7. Immediately prior to heat shock, add 36μL DMSO.

8. Heat shock was performed at 42 °C for 15 min.

9. The solution was spun down for 30 s, the liquids were removed, and re-suspend in 50μL $H_2O$.

10. Plated all 500μL on the selection plate to achieve the highest transformation efficiency.

**Supplemental method 2: Emulsion PCR set up with EURx Micellula DNA Emulsion & Purification (ePCR) PCR kit**

1. The emulsion-oil phase was prepared on ice.

Emulsion Oil Phase:

| Oil surfactant mixture (300 μl per reaction) | (μL) |
|---|---|
| Emulsion component 1 | 220 |
| Emulsion component 2 | 20 |
| Emulsion component 3 | 60 |

2. Mix thoroughly by vertexing at the highest level and put in a 4 °C cold room for further use.

3. Prepare the PCR water phase on ice ( check the additional setup for amplifying the TL region from the DNA pool at the end of this document).

PCR Water Phase - 1

| Emulsion PCR water phase using Kaplan lab Phusion for amplification from synthesized library pool | |
|---|---|
| Components | 1X |
| 10X dNTPs(μL) | 5 |
| 5X detergent free Buffer(μL) | 10 |
| $H_2O$(μL) | 32 |
| 100μM Primer F (μL) | 0.5 |
| 100μM Primer R (μL) | 0.5 |
| Agilent TL library (1ng/μL) | 1 |
| Phusion(μL) | 0.5 |
| 1mg/mL BSA | 0.5 |
| Total(μL) | 50 |

Additionally, PCR Water Phase – 2 for amplifying the TL region from extracted DNA pool

| For amplification from DNA pool | |
|---|---|
| Emulsified PCR using Kaplan Lab Phusion | |
| Components | |
| 10X dNTPs(μL) | 5 |
| 5X detergent free Buffer(μL) | 10 |
| $H_2O$(μL) | 32 |
| 100μM Primer F CKO413 (μL) | 0.5 |
| 100μM Primer R CKO414(μL) | 0.5 |
| Diluted 5ng/μL TL Lib6 DNA(μL) | 1 |
| Phusion(μL) | 0.5 |
| 1mg/mL BSA | 0.5 |
| Total(μL) | 50 |
| 16 Cycles for the reactions | |

4. Create emulsion reactions by mixing the 300μl precooled oil surfactant mixture and 50μl of precooled PCR water phase with vortexing at the maximum speed for 5 min in a cold room.
5. The solution was quickly spun down at ~1000rpm for 5 s. Dispense ~110μL aliquot into three PCR tubes.

6. PCR was performed according to the following protocol.
Note 1: Do not exceed the 95°C denaturing temperature because some buffers tend to destabilize the emulsion.
Note 2: The number of cycles was determined by Q-PCR with the same amount of template and oligos. A cycle was selected in the upper half of the linear amplification curve. Two more cycles were added to the selected cycle number because emulsion

PCR tends to have a lower amplification efficiency than the standard PCR. To confirm the linearity, a further test with selected cycle, selected cycle plus two more cycles, and selected cycle plus four more cycles was applied to three emulsion PCR reactions (from the same mix). The products of the three emulsion PCR were analyzed using agarose gel electrophoresis. The amount of PCR with the selected cycle plus two more cycles was normally between the other two cycles.
Example gel figure of three emulsion PCR reactions

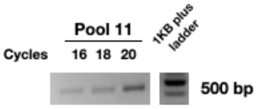


Thermal Cycling

| | | |
|---|---|---|
| 95 °C | 3min | |
| 95 °C | 15s | 18 cycles |
| 55 °C | 30s | |
| 72 °C | 30s/kb | |
| 72 °C | 5 min | |
| 12 °C | Forever | |

7. Once the PCR was completed, the corresponding triplicates of each ePCR assay were pooled into a single 2 ml reaction tube. Break emulsion by adding 1.0 ml 2-butanol (or butanol). Mix by vortexing.

8. Add 400 µL of orange-colored Orange-DX buffer to the opened emulsion solution. Mix the emulsion solution with gentle agitation (e.g., on a rotator for 2 min).

9. Centrifuge for 2 min at maximum speed (e.g. 16 000 x g / approx. 14 000 rpm) for phase separation.

10. Most of the yellow-colored organic phase was removed.

11. Apply 40 µL of activation Buffer DX onto the spin column (do not spin) and keep it at room temperature until the mixture is transferred to the spin column (at least 10 min).

12. Pour the mixture (aqueous phase + interphase; max. Six hundred microliters) into a spin column/receiver tube assembly.

13. Spin down in a microcentrifuge at 12,000 rpm for 1 min, discard the flow-through.

14. Add 500 µL of Wash-DX1 buffer and spin down at 12,000 rpm (~11.000 x g) for 1 min, and discard the flow-through.

15. Add 650 µL of Wash-DX2 buffer and spin down at 12,000 rpm (~11.000 × g) for 1 min, and discard the flow-through.

16. The mixture was spun down at 12,000 rpm (~11.000 x g) for 2 min to remove traces of Wash-DX buffer.

17. The spin column was placed into a new receiver tube (1.5-2 ml). Add 50-150 µl of Elution-DX buffer to elute the bound DNA.

18. Incubate the spin column/receiver tube assembly for 2 min at room temperature. Spin down at 12,000 rpm(~11.000 × g) for 1 min. The elution process was repeated.

**Supplemental Method 3: Amplification/transformation/screening of mutant libraries and sequencing pool preparation.**

I. Amplification of mutants from the synthesized pool.

1. The Agilent TL library was dissolved in 100μL of 10mM Tris buffer (pH8) as suggested by Cohen's protocol. 1μL of the library was used in each 50μL PCR reaction. The number of cycles was estimated using qPCR. Two rounds of emulsion PCR were performed to obtain sufficient library products. The size of the library products was confirmed by agarose gel electrophoresis.
2. Two flanking regions were added to the TL regions amplified by PCR sequencing, as shown in the following schematic.

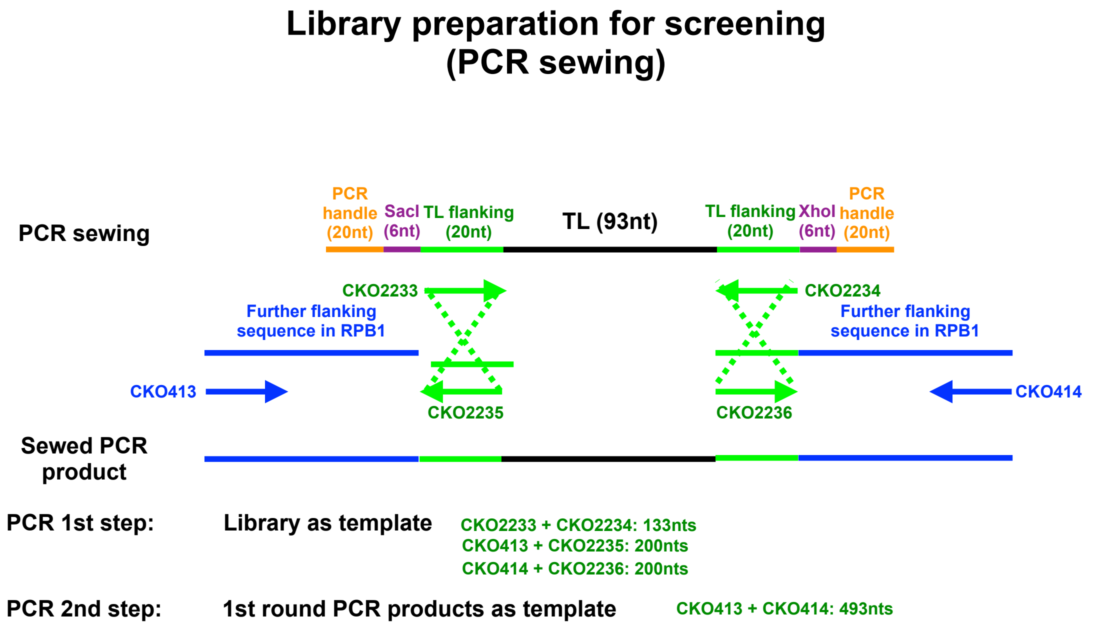


II. TL library screening.

1. Transformation was performed using a high-efficiency large-scale chemical yeast transformation protocol (Supplemental Method 1). 100ng of MluI digested plasmid and 383ng of variants library were used in the transformation.
2. Screening was performed using scraping and replating. The number of cells for each condition is listed in the following table.

| Condition | Cells plated | Days of collection |
|---|---|---|
| SC-Leu (Pre) | 100M | 2 days |
| SC-Leu + 5FOA | 400K | 3 days |
| SC-Leu (Post) | 400K | 2 days |
| SC-Lys | 100M | 8 days |

| | | |
|---|---|---|
| YPRaf | 400K | 4 days |
| YPRafGal | 50M | 7 days |
| SC-Leu + 20μg/mL MPA | 1M | 4 days |
| SC-Leu + 15mM Mn | 1M | 5 days |
| SC-Leu + 3% Formamide | 500K | 3 days |

III. Preparation of sequencing pool

We have 10 TL-screening mutant libraries that were tested under nine conditions; each library had three replicates, leading to 270 mutant pools (10 × 9 × 3 = 270).

1. DNA from mutant pools (n=270) was extracted using the Yeastar genomic DNA kit according to the manufacturer's instructions (Zymo Research), except that we used 1-5 x $10^8$ cells.
2. To amplify the TL region from the extracted DNA and add barcodes to distinguish 10 libraries, the TL regions of 270 mutant pools were amplified by standard PCR with 10 pairs of barcoded primers (one pair of primers with a certain barcode was used for all conditions and replicates of that library). PCR cycles were determined by Q-PCR to minimize the allele frequency shifts caused by amplification. We did Q-PCR test with two or three replicates of each library to determine a cycle that was in the linear range of the amplification curve. All three replicates of each library were subjected to the same cycle, which represents the average value of the replicates in the Q-PCR test. The determined cycles are shown in the following figure.

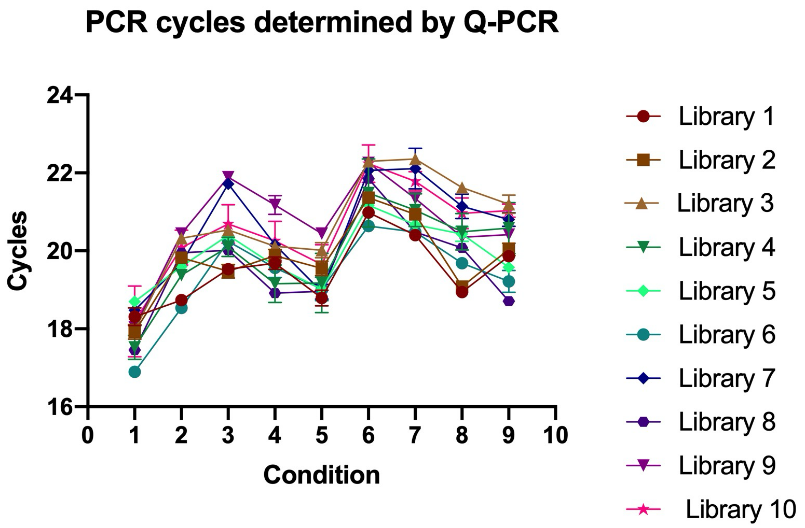


3. PCR products from step 2, representing libraries screened under the same conditions, were pooled. We obtained 27 pools in total after a combination of 27 conditions. To limit template switching, these 27 pools were amplified using emulsion PCR technology (EURx Micellula DNA Emulsion & Purification (ePCR PCR kit). Q-PCR was performed to determine the amplification cycle that was in the linear range for each pool. NEB barcodes containing primers were used to distinguish different conditions (NEBNext Multiplex Oligos for Illumina).

After two rounds of amplification, a sample-specific barcode sequence was added to the TL variants, and an adequate amount of TL variants was ready for sequencing. The indexed pooled samples were sequenced by single-end sequencing on an Illumina Next-seq (150nt reads). On average, over 11 million reads were obtained for individual samples with high reproducibility after two rounds of sequencing.

**Supplemental Method 4: Formulas of calculating functional interactions.**

For multiple mutant $M_1M_2M_3$ with count $c(M_1M_2M_3)$, we computed the ratio $rc(M_1M_2M_3) = c(M_1M_2M_3)/c(WT)$.

(1) The observed fitness of the multiple mutant $M_1M_2M_3$ is:

$$\log\left(\frac{rc\ (M1M2M3)sele}{rc(M1M2M3)unsele}\right)$$

(2) The expected fitness of a mutant $M_1M_2M_3$ is the log additive of the constituent single mutants $M_1$, $M_2$, and $M_3$.

$$\log\left(\frac{rc\ (M1)sele}{rc(M1)unsele}\right) + \log\left(\frac{rc(M2)sele}{rc(M2)unsele}\right) + \log\left(\frac{rc\ (M3)sele}{rc(M3)unsele}\right) = \log\left(\frac{rc(M1)sele * rc\ (M2)sele * rc\ (M3)sele}{rc\ (M1)unele * rc\ (M2)unsele * rc\ (M3)unsele}\right)$$

(3) We compared the fitness of $M_1M_2M_3$ with the log sum of its constituent $M_1$, $M_2$, and $M_3$ (compare the observed to the expected fitness), which is

$$Deviation\ score\ = \log\left(\frac{rc\ (M1M2M3)sele}{rc(M1M2M3)unsele}\right) - \log\left(\frac{rc(M1)sele * rc\ (M2)sele * rc\ (M3)sele}{rc\ (M1)unele * rc\ (M2)unsele * rc\ (M3)unsele}\right)$$

$$= \log\left(\frac{rc\ (M1M2M3)sele}{rc(M1)sele * rc\ (M2)sele * rc\ (M3)sele}\right) - \log\left(\frac{rc(M1M2M3)unsele}{rc\ (M1)unele * rc\ (M2)unsele * rc\ (M3)unsele}\right)$$

If

$$-1 < \log\left(\frac{rc\ (M1M2M3)sele}{rc(M1)sele * rc\ (M2)sele * rc\ (M3)sele}\right) - \log\left(\frac{rc(M1M2M3)unsele}{rc\ (M1)unele * rc\ (M2)unsele * rc\ (M3)unsele}\right) < 1$$

Then interaction among the constituent single mutants is additive.

If

$$\log\left(\frac{rc\ (M1M2M3)sele}{rc(M1)sele * rc\ (M2)sele * rc\ (M3)sele}\right) - \log\left(\frac{rc(M1M2M3)unsele}{rc\ (M1)unele * rc\ (M2)unsele * rc\ (M3)unsele}\right) \geq 1$$

Then the interaction is non-additive, positive interactions, including suppression and epistasis.

If

$$\log\left(\frac{rc\ (M1M2M3)sele}{rc(M1)sele * rc\ (M2)sele * rc\ (M3)sele}\right) - \log\left(\frac{rc(M1M2M3)unsele}{rc\ (M1)unele * rc\ (M2)unsele * rc\ (M3)unsele}\right) \leq -1$$

Then the interaction is non-additive, negative interactions, including synthetic sick, synthetic lethal and sign epistasis.

1. Gietz, R.D. and R.H. Schiestl, *High-efficiency yeast transformation using the LiAc/SS carrier DNA/PEG method.* Nat Protoc, 2007. **2**(1): p. 31-4.
