## Supplemental Table 1 for "Widespread epistasis shapes RNA Polymerase II active site function and evolution"

| Index | Handled library | WT | Allele number |
|---|---|---|---|
| **TL Lib1** | **Singles** | **111** | 620 |
| **TL Lib2** | **Pairwise Doubles** | **700** | 3914 |
| **TL Lib3** | **Evo present** | **123** | 662 |
| **TL Lib4** | **Evo Path** | **356** | 1987 |
| **TL Lib5** | **Coupling** | **130** | 724 |
| **TL Lib6** | **Target Doubles** | **858** | 4800 |
| **TL Lib7** | **T834P-target** | **111** | 621 |
| **TL Lib8** | **T834A-target** | **111** | 621 |
| **TL Lib9** | **Y769F-target** | **111** | 621 |
| **TL Lib10** | **S713P-target** | **111** | 621 |
| **Total** | | **2722** | 15191 |
