## Supplemental Table 2 for "Widespread epistasis shapes RNA Polymerase II active site function and evolution"

| single 1/single2 | Plasmid | Ref | Derivation of the mutants | Spot growth defect | spot interaction | High thorughput growth defect | Deviation score | High thorughput interaction | Consistent? |
|---|---|---|---|---|---|---|---|---|---|
| N1082S/E1103G | pCK898 | [1] | Site-directed mutagenesis | Viable | Suppression | -0.107717799 | 6.392282201 | Suppression | Consistent |
| N1082A/G1097D | pCK1009 | [1] | Site-directed mutagenesis | Viable | Suppression | -0.609228746 | 5.890771254 | Suppression | Consistent |
| Q1078S/E1103G | pCK896 | [1] | Site-directed mutagenesis | Viable | Suppression | -0.72462063 | 5.77537937 | Suppression | Consistent |
| T834A/E1103G | pCK1410 | [2] | Site-directed mutagenesis | Viable | Suppression | -0.916984546 | 5.583015454 | Suppression | Consistent |
| N1082A/E1103G | pCK897 | [1] | Site-directed mutagenesis | Viable | Suppression | -1.398533462 | 5.101466538 | Suppression | Consistent |
| H1085Y/E1103G | pCK890 | [1] | Site-directed mutagenesis | Viable | Suppression | -1.84909122 | 4.65090878 | Suppression | Consistent |
| H1085Y/E1103G | pCK890 | [1] | Site-directed mutagenesis | Viable | Suppression | -1.84909122 | 4.65090878 | Suppression | Consistent |
| H1085Q/E1103G | pCK901 | [1] | Site-directed mutagenesis | Viable | Suppression | -1.849515981 | 4.650484019 | Suppression | Consistent |
| Q1078A/G1097D | pCK1008 | [1] | Site-directed mutagenesis | Viable | Suppression | -1.88786578 | 4.61213422 | Suppression | Consistent |
| F1086S/E1103G | pCK914 | [1] | Site-directed mutagenesis | Viable | Suppression | -1.490457626 | 4.292969551 | Suppression | Consistent |
| Q1078A/E1103G | pCK947 | [1] | Site-directed mutagenesis | Viable | Suppression | -2.691382561 | 3.808617439 | Suppression | Consistent |
| H1085R/E1103G | pCK902 | [1] | Site-directed mutagenesis | Viable | Suppression | -3.687573729 | 2.812426271 | Suppression | Consistent |
| H1085A/G1097D | pCK1007 | [1] | Site-directed mutagenesis | Viable | Suppression | -4.097056362 | 2.402943638 | Suppression | Consistent |
| H1085A/E1103G | pCK899 | [1] | Site-directed mutagenesis | Viable | Suppression | -4.166242681 | 2.333757319 | Suppression | Consistent |
| L1081F/E1103G | pCK939 | [1] | Site-directed mutagenesis | Viable | Suppression | -5.100889826 | 1.399110174 | Suppression | Consistent |
| H1085W/E1103G | pCK903 | [1] | Site-directed mutagenesis | Viable | Suppression | -5.189712459 | 1.310287541 | Suppression | Consistent |
| T834P/N1082S | pCK1894 | [2] | Site-directed mutagenesis | Viable | Suppression | -5.431195104 | 1.068804896 | Suppression | Consistent |
| Q1078S/F1084I | pCK1184 | [1] | Site-directed mutagenesis | Viable | Suppression of Q1078S | -5.75051526 | 0.74948474 | Suppression of Q1078S | Consistent |
| N1082S/F1084I | pCK951 | [1] | Site-directed mutagenesis | Viable | Synthetic sickness or additive | -6.165664446 | 0.334335554 | Additive | Consistent |
| F1084I/E1103G | pCK952 | [1] | Site-directed mutagenesis | Inviable | Double mutant lethality | ultra sick/Lethal | Double mutant lethality | Double mutant sickness or lethality explainable by synthetic interaction | Consistent |
| H1085Y/F1086S | pCK894 | [1] | Site-directed mutagenesis | Inviable | Double mutant lethality | ultra sick/Lethal | Double mutant lethality | Double mutant sickness or lethality explainable by synthetic interaction | Consistent |
| H1085Y/F1086Y | pCK893 | [1] | Site-directed mutagenesis | Inviable | Double mutant lethality | ultra sick/Lethal | Double mutant lethality | Double mutant sickness or lethality explainable by synthetic interaction | Consistent |
| N1082R/H1085Y | pCK877 | [1] | Site-directed mutagenesis | Inviable | Double mutant lethality | ultra sick/Lethal | Double mutant lethality | Double mutant sickness or lethality explainable by synthetic interaction | Consistent |
| T834A/N1082S | pCK1401 | [2] | Site-directed mutagenesis | Inviable | Double mutant lethality | ultra sick/Lethal | Double mutant lethality | Double mutant sickness or lethality explainable by synthetic interaction | Consistent |
| T834A/H1085Q | pCK1402 | [2] | Site-directed mutagenesis | Inviable | Double mutant lethality | ultra sick/Lethal | Double mutant lethality | Double mutant sickness or lethality explainable by synthetic interaction | Consistent |
| T834A/F1086S | pCK1407 | [2] | Site-directed mutagenesis | Inviable | Double mutant lethality | ultra sick/Lethal | Double mutant lethality | Double mutant sickness or lethality explainable by synthetic interaction | Consistent |
| T834P/F1084I | pCK1115 | [2] | Site-directed mutagenesis | Inviable | Double mutant lethality | ultra sick/Lethal | Double mutant lethality | Double mutant sickness or lethality explainable by synthetic interaction | Consistent |
| T834P/E1103G | pCK1116 | [2] | Site-directed mutagenesis | Inviable | Double mutant lethality | ultra sick/Lethal | Double mutant lethality | Double mutant sickness or lethality explainable by synthetic interaction | Consistent |
| G1097D/E1103G | pCK1189 | [1] | Site-directed mutagenesis | Inviable | Double mutant lethality | ultra sick/Lethal | Double mutant lethality | Double mutant sickness or lethality explainable by synthetic interaction | Consistent |
| F1084I/H1085Y | pCK876 | [1] | Site-directed mutagenesis | Inviable | Double mutant lethality | ultra sick/Lethal | Double mutant lethality | Double mutant lethality explainable by presence of lethal single mutant | Consistent |
| L1081A/E1103G | pCK948 | [1] | Site-directed mutagenesis | Inviable | Double mutant lethality | ultra sick/Lethal | Double mutant lethality | Double mutant lethality explainable by presence of lethal single mutant | Consistent |
| L1081G/E1103G | pCK936 | [1] | Site-directed mutagenesis | Inviable | Double mutant lethality | ultra sick/Lethal | Double mutant lethality | Double mutant lethality explainable by presence of lethal single mutant | Consistent |
| Q1078A/H1085A | pCK950 | [1] | Site-directed mutagenesis | Inviable | Double mutant lethality | ultra sick/Lethal | Double mutant lethality | Double mutant lethality explainable by presence of lethal single mutant | Consistent |
| Q1078A/H1085Y | pCK940 | [1] | Site-directed mutagenesis | Inviable | Double mutant lethality | ultra sick/Lethal | Double mutant lethality | Double mutant lethality explainable by presence of lethal single mutant | Consistent |
| Q1078A/N1082A | pCK945 | [1] | Site-directed mutagenesis | Inviable | Double mutant lethality | ultra sick/Lethal | Double mutant lethality | Double mutant lethality explainable by presence of lethal single mutant | Consistent |
| Q1078S/H1085A | pCK919 | [1] | Site-directed mutagenesis | Inviable | Double mutant lethality | ultra sick/Lethal | Double mutant lethality | Double mutant lethality explainable by presence of lethal single mutant | Consistent |
| T834A/Q1078A | pCK1403 | [2] | Site-directed mutagenesis | Inviable | Double mutant lethality | ultra sick/Lethal | Double mutant lethality | Double mutant lethality explainable by presence of lethal single mutant | Consistent |
| T834A/H1085A | pCK1399 | [2] | Site-directed mutagenesis | Inviable | Double mutant lethality | ultra sick/Lethal | Double mutant lethality | Double mutant lethality explainable by presence of lethal single mutant | Consistent |
| T834A/N1082A | pCK1411 | [2] | Site-directed mutagenesis | Inviable | Double mutant lethality | ultra sick/Lethal | Double mutant lethality | Double mutant lethality explainable by presence of lethal single mutant | Consistent |
| T834A/H1085Y | pCK1408 | [2] | Site-directed mutagenesis | Inviable | Double mutant lethality | ultra sick/Lethal | Double mutant lethality | Double mutant lethality explainable by presence of lethal single mutant | Consistent |
| T834A/Q1078S | pCK1400 | [2] | Site-directed mutagenesis | Inviable | Double mutant lethality | ultra sick/Lethal | Double mutant lethality | Double mutant lethality explainable by presence of lethal single mutant | Consistent |
| T834P/Q1078A | pCK1890 | [2] | Site-directed mutagenesis | Inviable | Double mutant lethality | ultra sick/Lethal | Double mutant lethality | Double mutant lethality explainable by presence of lethal single mutant | Consistent |
| T834P/H1085A | pCK1891 | [2] | Site-directed mutagenesis | Inviable | Double mutant lethality | ultra sick/Lethal | Double mutant lethality | Double mutant lethality explainable by presence of lethal single mutant | Consistent |
| T834P/N1082A | pCK1892 | [2] | Site-directed mutagenesis | Inviable | Double mutant lethality | ultra sick/Lethal | Double mutant lethality | Double mutant lethality explainable by presence of lethal single mutant | Consistent |
| N1082A/H1085Y | pCK917 | [1] | Site-directed mutagenesis | Inviable | Double mutant lethality | ultra sick/Lethal | Double mutant lethality | Double mutant lethality explainable by presence of lethal single mutant | Consistent |
| N1082S/H1085Y | pCK1185 | [1] | Site-directed mutagenesis | Inviable | Double mutant lethality | ultra sick/Lethal | Double mutant lethality | Double mutant lethality explainable by presence of lethal single mutant | Consistent |
| Q1078S/H1085Y | pCK918 | [1] | Site-directed mutagenesis | Inviable | Double mutant lethality | ultra sick/Lethal | Double mutant lethality | Double mutant lethality explainable by presence of lethal single mutant | Consistent |
| Q1078S/H1085Q | pCK1183 | [1] | Site-directed mutagenesis | Inviable | Double mutant lethality | ultra sick/Lethal | Double mutant lethality | Double mutant lethality explainable by synthetic lethality | Consistent |
| N1082A/F1084I | pCK954 | [1] | Site-directed mutagenesis | Viable | Suppression | ultra sick/Lethal | Double mutant lethality | Double mutant lethality explainable by presence of lethal single mutant | Inconsistent due to mutants in ultra sick/lethal range |
| H1085F/E1103G | pCK900 | [1] | Site-directed mutagenesis | Viable | Suppression | ultra sick/Lethal | Double mutant lethality | Double mutant sickness or lethality explainable by synthetic interaction | Inconsistent due to mutants in ultra sick/lethal range |
| N1082R/F1084I | pCK875 | [1] | Site-directed mutagenesis | Viable | Suppression | ultra sick/Lethal | Double mutant lethality | Double mutant sickness or lethality explainable by synthetic interaction | Inconsistent due to mutants in ultra sick/lethal range |
| Q1078N/E1103G | pCK895 | [1] | Site-directed mutagenesis | Viable | Suppression | ultra sick/Lethal | Double mutant lethality | Double mutant lethality explainable by presence of lethal single mutant | Inconsistent due to mutants in ultra sick/lethal range |
| T834P/H1085Y | pCK1113 | [2] | Site-directed mutagenesis | Viable | Suppression | ultra sick/Lethal | Double mutant lethality | Double mutant lethality explainable by presence of lethal single mutant | Inconsistent due to mutants in ultra sick/lethal range |
| T834P/Q1078S | pCK1893 | [2] | Site-directed mutagenesis | Viable | Suppression | ultra sick/Lethal | Double mutant lethality | Double mutant lethality explainable by presence of lethal single mutant | Inconsistent due to mutants in ultra sick/lethal range |
| Q1078S/N1082A | pCK904 | [1] | Site-directed mutagenesis | Viable | Suppression | ultra sick/Lethal | Double mutant lethality | Double mutant lethality explainable by presence of lethal single mutant | Inconsistent due to mutants in ultra sick/lethal range |
| T834P/F1086S | pCK1114 | [2] | Site-directed mutagenesis | Viable | Conditional suppression or enhancement | ultra sick/Lethal | Double mutant lethality | Double mutantsickness or lethality explainable by synthetic interaction | Inconsistent due to mutants in ultra sick/lethal range |
| Q1078S/N1082S | pCK1186 | [1] | Site-directed mutagenesis | Viable | Epistasis | ultra sick/Lethal | Double mutant lethality | Double mutant lethality explainable by presence of lethal single mutant | Inconsistent due to mutants in ultra sick/lethal range |
| T834P/H1085Q | pCK1895 | [2] | Site-directed mutagenesis | Viable | Sick combination | ultra sick/Lethal | Double mutant lethality | Double mutant sickness or lethality explainable by synthetic interaction | Inconsistent due to mutants in ultra sick/lethal range |
| F1084I/H1085Q | pCK953 | [1] | Site-directed mutagenesis | Viable | Synthetic sickness | ultra sick/Lethal | Double mutant lethality | Double mutant sickness or lethality explainable by synthetic interaction | Inconsistent due to mutants in ultra sick/lethal range |


function and Pol II activity-dependent control of start site selection in vivo. PLoS Genet 8, e1002627 (2012).

er, J. Van den Brulle, S. H. Sze, C. D. Kaplan, High-Resolution Phenotypic Landscape of the RNA Polymerase II Trigger Loop. *PLoS Genet* **12**, e1006321 (2016).
