## Supplemental Table 3 for "Widespread epistasis shapes RNA Polymerase II active site function and evolution"

| Name | Genotype | Note |
| --- | --- | --- |
| pCK892 | LEU2 CEN ARS ampr ColE1 ori rpb1 ∆Trigger Loop,T69 corrected | WT *RPB1* plasmid with TL deleted |
| pCK2193 | LEU2 CEN ARS ampr ColE1 ori rpb1 T834P ∆Trigger Loop | T834P plasmid with TL deleted |
| pCK2194 | LEU2 CEN ARS ampr ColE1 ori rpb1 T834A ∆Trigger Loop | T834A plasmid with TL deleted |
| pCK2198 | LEU2 CEN ARS ampr ColE1 ori rpb1 S713P ∆Trigger Loop | S713P plasmid with TL deleted |
| CKY283 | ura3-52 his3∆200 leu2∆1 or ∆0 trp1∆63 met15∆0 lys2-128∂ gal10∆56 rpb1∆::CLONATMX RPB3::TAP::KlacTRP1 | WT yeast strain |
| CKY3208 | ura3-52 his3∆200 leu2∆1 or ∆0 trp1∆63 met15∆0 lys2-128∂ gal10∆56 rpb1∆::CLONATMX RPB3::TAP::KlacTRP1 rpb2 Y769F | *rpb2* Y769F strain |
