## Supplemental Table 4 for "Widespread epistasis shapes RNA Polymerase II active site function and evolution"

| Name | Sequence | Description | Amplification | Schematic | Note |
|---|---|---|---|---|---|
| **CKO2751** | ACACTCTTTCCCTACACGACGCTCTTCCGATCTGTCGGTAATTAGCAGCCCAATCCATTGGTG | TL Library amplification (First amplification) | TL library 1 | R1-BC1-N-TL-F | Custom primer |
| **CKO2752** | ACACTCTTTCCCTACACGACGCTCTTCCGATCTAGGTCACTAGTAGCAGCCCAATCCATTGGTG | TL Library amplification (First amplification) | TL library 2 | R1-BC2-NN-TL-F | Custom primer |
| **CKO2753** | ACACTCTTTCCCTACACGACGCTCTTCCGATCTGAATCCGACACTAGCAGCCCAATCCATTGGTG | TL Library amplification (First amplification) | TL library 3 | R1-BC3-NNN-TL-F | Custom primer |
| **CKO2754** | ACACTCTTTCCCTACACGACGCTCTTCCGATCTGTACCTTGGCTCTAGCAGCCCAATCCATTGGTG | TL Library amplification (First amplification) | TL library 4 | R1-BC4-NNNN-TL-F | Custom primer |
| **CKO2755** | ACACTCTTTCCCTACACGACGCTCTTCCGATCTCATGAGGATTCGCTAGCAGCCCAATCCATTGGTG | TL Library amplification (First amplification) | TL library 5 | R1-BC5-NNNNN-TL-F | Custom primer |
| **CKO2756** | ACACTCTTTCCCTACACGACGCTCTTCCGATCTTGACTGACACGAACTAGCAGCCCAATCCATTGGTG | TL Library amplification (First amplification) | TL library 6 | R1-BC6-NNNNNN-TL-F | Custom primer |
| **CKO2757** | ACACTCTTTCCCTACACGACGCTCTTCCGATCTTCAGACGAGAGTTGTTAGCAGCCCAATCCATTGGTG | TL Library amplification (First amplification) | TL library 7 | R1-BC7-NNNNNNN-TL-F | Custom primer |
| **CKO2758** | ACACTCTTTCCCTACACGACGCTCTTCCGATCTGATAGGCTCTCTGTGTTAGCAGCCCAATCCATTGGTG | TL Library amplification (First amplification) | TL library 8 | R1-BC8-NNNNNNNN-TL-F | Custom primer |
| **CKO2759** | ACACTCTTTCCCTACACGACGCTCTTCCGATCTTGGTACAGTGTGCTCCTTAGCAGCCCAATCCATTGGTG | TL Library amplification (First amplification) | TL library 9 | R1-BC9-NNNNNNNNN-TL-F | Custom primer |
| **CKO2760** | ACACTCTTTCCCTACACGACGCTCTTCCGATCTCAAGGTCTGGACAGTTATTAGCAGCCCAATCCATTGGTG | TL Library amplification (First amplification) | TL library 10 | R1-BC10-NNNNNNNNNN-TL-F | Custom primer |
| **CKO2761** | GTGACTGGAGTTCAGACGTGTGCTCTTCCGATCTGAAGGGGTTTTCATGTTTTTGG | TL Library amplification (First amplification) | TL libraries | R2-TL-R | Custom primer |
| P17-B5 | AGGATAGC | NEB index oligos (Second amplification) | Condition 1-1 | | NEB index oligos |
| P18-B6 | CCTTCCAT | NEB index oligos (Second amplification) | Condition 1-2 | | NEB index oligos |
| P19-B7 | GTCCTTGA | NEB index oligos (Second amplification) | Condition 1-3 | | NEB index oligos |
| P20-B8 | TGCGTAAC | NEB index oligos (Second amplification) | Condition 2-1 | | NEB index oligos |
| P21-B9 | CACAGACT | NEB index oligos (Second amplification) | Condition 2-2 | | NEB index oligos |
| P22-B10 | TTACGTGC | NEB index oligos (Second amplification) | Condition 2-3 | | NEB index oligos |
| P23-B11 | CCAAGGTT | NEB index oligos (Second amplification) | Condition 3-1 | | NEB index oligos |
| P24-B12 | CACGCAAT | NEB index oligos (Second amplification) | Condition 3-2 | | NEB index oligos |
| P53-E5 | CCGCTTAA | NEB index oligos (Second amplification) | Condition 3-3 | | NEB index oligos |
| P54-E6 | TACCTGCA | NEB index oligos (Second amplification) | Condition 4-1 | | NEB index oligos |
| P55-E7 | GTCGATTG | NEB index oligos (Second amplification) | Condition 4-2 | | NEB index oligos |
| P56-E8 | TATGGCAC | NEB index oligos (Second amplification) | Condition 4-3 | | NEB index oligos |
| P57-E9 | CTCGAACA | NEB index oligos (Second amplification) | Condition 5-1 | | NEB index oligos |
| P58-E10 | CAACTCCA | NEB index oligos (Second amplification) | Condition 5-2 | | NEB index oligos |
| P59-E11 | GTCATCGT | NEB index oligos (Second amplification) | Condition 5-3 | | NEB index oligos |
| P60-E12 | GGACATCA | NEB index oligos (Second amplification) | Condition 6-1 | | NEB index oligos |
| P85-H1 | TACTCCAG | NEB index oligos (Second amplification) | Condition 6-2 | | NEB index oligos |
| P86-H2 | GGAAGAGA | NEB index oligos (Second amplification) | Condition 6-3 | | NEB index oligos |
| P87-H3 | GCGTTAGA | NEB index oligos (Second amplification) | Condition 7-1 | | NEB index oligos |
| P88-H4 | ATCTGACC | NEB index oligos (Second amplification) | Condition 7-2 | | NEB index oligos |
| P89-H5 | AACCAGAG | NEB index oligos (Second amplification) | Condition 7-3 | | NEB index oligos |
| P90-H6 | GTACCACA | NEB index oligos (Second amplification) | Condition 8-1 | | NEB index oligos |
| P91-H7 | GGTATAGG | NEB index oligos (Second amplification) | Condition 8-2 | | NEB index oligos |
| P92-H8 | CGAGAGAA | NEB index oligos (Second amplification) | Condition 8-3 | | NEB index oligos |
| P93-H9 | CAGCATAC | NEB index oligos (Second amplification) | Condition 9-1 | | NEB index oligos |
| P94-H10 | CTCGACTT | NEB index oligos (Second amplification) | Condition 9-2 | | NEB index oligos |
| P95-H11 | CTTCGGTT | NEB index oligos (Second amplification) | Condition 9-3 | | NEB index oligos |
| P96-H12 | CCACAACA | NEB index oligos (Second amplification) | Condition 0 | | NEB index oligos |
