## Supplemental Table 5 for "Widespread epistasis shapes RNA Polymerase II active site function and evolution"

| Condition | Cells plated | Days for phenotyping |
|---|---|---|
| SC-Leu (Pre) | 100M | 2 days |
| SC-Leu + 5FOA | 400K | 3 days |
| SC-Leu (Post) | 400K | 2 days |
| SC-Lys | 100M | 7 days |
| YPRaf | 400K | 6 days |
| YPRafGal | 50M | 6 days |
| SC-Leu + 20μg/mL MPA | 1M | 4 days |
| SC-Leu + 15mM Mn | 1M | 5 days |
| SC-Leu + 3% Formamide | 500K | 3 days |
