## Supplemental Table 6 for "Widespread epistasis shapes RNA Polymerase II active site function and evolution"

### GOF

Regression type: Logistic regression

| Model | | | | |
|---|---|---|---|---|
| Parameter estimates | Variable | Estimate | Standard | 95% CI (profile likelihood) |
| β0 | Intercept | -1.816 | 1.549 | -5.879 to 0.8657 |
| β1 | MPA | -2.542 | 1.294 | -6.196 to -0.6109 |
| β2 | Lys | 1.942 | 1.121 | 0.4256 to 5.021 |
| β3 | Gal | -0.0657 | 0.282 | -0.7183 to 0.5200 |
| β4 | MPA : Lys | 0.5297 | 0.2843 | -0.2079 to 1.322 |
| β5 | MPA : Gal | 0.08373 | 0.276 | -0.5768 to 0.5721 |
| β6 | Lys : Gal | -0.0256 | 0.2288 | -0.2720 to 0.5072 |

| Odds ratios | Variable | Estimate | 95% CI (profile likelihood) |
|---|---|---|---|
| β0 | Intercept | 0.1626 | 0.002798 to 2.377 |
| β1 | MPA | 0.07874 | 0.002037 to 0.5429 |
| β2 | Lys | 6.974 | 1.530 to 151.6 |
| β3 | Gal | 0.9364 | 0.4876 to 1.682 |
| β4 | MPA : Lys | 1.698 | 0.8123 to 3.752 |
| β5 | MPA : Gal | 1.087 | 0.5617 to 1.772 |
| β6 | Lys : Gal | 0.9748 | 0.7618 to 1.661 |

| Model diagnostics | Degrees of Freedom | AICc |
|---|---|---|
| Intercept-only model | 60 | 84.64 |
| Selected model | 54 | 31.1 |

| Area under the ROC curve | |
|---|---|
| Area | 0.9889 |
| Std. Error | 0.009628 |
| 95% confidence interval | 0.9700 to 1.000 |
| P value | <0.0001 |

| Classification table | Predicted Not GOF | Predicted | Total | % Correctly classified |
|---|---|---|---|---|
| Observed Not GOF | 36 | 0 | 36 | 100 |
| Observed GOF | 2 | 23 | 25 | 92 |
| Total | 38 | 23 | 61 | 96.72 |

| | |
|---|---|
| Negative predictive power (%) | 94.74 |
| Positive predictive power (%) | 100 |

| | |
|---|---|
| Classification cutoff | 0.75 |

| Pseudo R squared | |
|---|---|
| Tjur's R squared | 0.8463 |

| Data summary | |
|---|---|
| Rows in table | 65 |
| Rows skipped (missing data) | 4 |
| Rows analyzed (#observations) | 61 |
| Number of GOF | 25 |
| Number of Not GOF | 36 |
| Number of parameter estimates | 7 |
| #observations/#parameters | 8.7 |
| # of GOF/#parameters | 3.6 |
| # of Not GOF/#parameters | 5.1 |

### LOF

Regression type: Logistic regression

| Model | | | | |
|---|---|---|---|---|
| Parameter estimates | Variable | Estimate | Standard | 95% CI (profile likelihood) |
| β0 | Intercept | -1.916 | 1.327 | -5.389 to 0.3409 |
| β1 | MPA | 1.392 | 1.508 | -0.7599 to 5.252 |
| β2 | Spt | 1.328 | 0.9944 | -0.04761 to 4.209 |
| β3 | Gal | 0.8353 | 0.4913 | 0.1660 to 2.396 |
| β4 | MPA : Spt | 0.01112 | 0.1707 | -0.3174 to 0.5524 |
| β5 | MPA : Gal | 0.2992 | 0.3107 | -0.2797 to 1.150 |
| β6 | Spt : Gal | -0.8823 | 0.6035 | -2.785 to -0.1533 |

| Odds ratios | Variable | Estimate | 95% CI (profile likelihood) |
|---|---|---|---|
| β0 | Intercept | 0.1472 | 0.004566 to 1.406 |
| β1 | MPA | 4.021 | 0.4677 to 191.0 |
| β2 | Spt | 3.774 | 0.9535 to 67.29 |
| β3 | Gal | 2.305 | 1.181 to 10.98 |
| β4 | MPA : Spt | 1.011 | 0.7281 to 1.737 |
| β5 | MPA : Gal | 1.349 | 0.7560 to 3.159 |
| β6 | Spt : Gal | 0.4138 | 0.06175 to 0.8579 |

| Model diagnostics | Degrees of Freedom | AICc |
|---|---|---|
| Intercept-only model | 60 | 86.48 |
| Selected model | 54 | 29.93 |

| Area under the ROC curve | |
|---|---|
| Area | 0.9914 |
| Std. Error | 0.007495 |
| 95% confidence interval | 0.9767 to 1.000 |
| P value | <0.0001 |

| Classification table | Predicted 0 | Predicted | Total | % Correctly classified |
|---|---|---|---|---|
| Observed 0 | 32 | 0 | 32 | 100 |
| Observed 1 | 3 | 26 | 29 | 89.66 |
| Total | 35 | 26 | 61 | 95.08 |

| | |
|---|---|
| Negative predictive power (%) | 91.43 |
| Positive predictive power (%) | 100 |

| | |
|---|---|
| Classification cutoff | 0.75 |

| Pseudo R squared | |
|---|---|
| Tjur's R squared | 0.8505 |

| Data summary | |
|---|---|
| Rows in table | 65 |
| Rows skipped (missing data) | 4 |
| Rows analyzed (#observations | 61 |
| Number of 1 | 29 |
| Number of 0 | 32 |
| Number of parameter estimate | 7 |
| #observations/#parameters | 8.7 |
| # of 1/#parameters | 4.1 |
| # of 0/#parameters | 4.6 |
