## Supplemental Table 7 for "Widespread epistasis shapes RNA Polymerase II active site function and evolution"

| **Kruskal-Wallis test with deviation scores of probe mutants when they were combined with viable mutants** | | | | |
|---|---|---|---|---|
| Dunn's multiple comparisons t | Mean rank diff. | Significant? | Summary | Adjusted P Value |
| S713P vs. Y769F | 118.1 | No | ns | 0.8279 |
| S713P vs. E1103G | 7.282 | No | ns | >0.9999 |
| S713P vs. L1101S | 270.8 | Yes | **** | <0.0001 |
| S713P vs. F1084I | 383 | Yes | **** | <0.0001 |
| S713P vs. M1079V | -62.19 | No | ns | >0.9999 |
| S713P vs. T834P | 610.7 | Yes | **** | <0.0001 |
| Y769F vs. E1103G | -110.8 | No | ns | >0.9999 |
| Y769F vs. L1101S | 152.7 | No | ns | 0.1754 |
| Y769F vs. F1084I | 264.9 | Yes | **** | <0.0001 |
| Y769F vs. M1079V | -180.3 | Yes | * | 0.0393 |
| Y769F vs. T834P | 492.6 | Yes | **** | <0.0001 |
| E1103G vs. L1101S | 263.5 | Yes | *** | 0.0001 |
| E1103G vs. F1084I | 375.7 | Yes | **** | <0.0001 |
| E1103G vs. M1079V | -69.47 | No | ns | >0.9999 |
| E1103G vs. T834P | 603.4 | Yes | **** | <0.0001 |
| L1101S vs. F1084I | 112.2 | No | ns | >0.9999 |
| L1101S vs. M1079V | -333 | Yes | **** | <0.0001 |
| L1101S vs. T834P | 339.9 | Yes | **** | <0.0001 |
| F1084I vs. M1079V | -445.2 | Yes | **** | <0.0001 |
| F1084I vs. T834P | 227.7 | Yes | ** | 0.0017 |
| M1079V vs. T834P | 672.9 | Yes | **** | <0.0001 |
| Dunn's multiple comparisons t | Mean rank diff. | Significant? | Summary | Adjusted P Value |
| H1085L vs. H1085Y | 236.3 | Yes | **** | <0.0001 |
| H1085L vs. N1082S | 6.459 | No | ns | >0.9999 |
| H1085L vs. Q1078S | 244.4 | Yes | **** | <0.0001 |
| H1085L vs. T834A | -25.35 | No | ns | >0.9999 |
| H1085Y vs. N1082S | -229.8 | Yes | **** | <0.0001 |
| H1085Y vs. Q1078S | 8.097 | No | ns | >0.9999 |
| H1085Y vs. T834A | -261.6 | Yes | **** | <0.0001 |
| N1082S vs. Q1078S | 237.9 | Yes | **** | <0.0001 |
| N1082S vs. T834A | -31.8 | No | ns | >0.9999 |
| Q1078S vs. T834A | -269.7 | Yes | **** | <0.0001 |
